# Identification of a Highly Expressed Gene Cluster Likely Coding for Benzene Activation Enzymes in a Methanogenic Enrichment Culture

**DOI:** 10.1101/2024.12.15.628547

**Authors:** Courtney R. A. Toth, Olivia Molenda, Camilla Nesbø, Fei Luo, Cheryl E. Devine, Xu Chen, Kan Wu, Johnny Z. Xiao, Rishika Puri, Shen Guo, Nancy Bawa, Po-Hsiang Wang, Yifeng Wei, Robert Flick, Elizabeth A. Edwards

**Author notes:** Liven Proteins Corporation, Mississauga, Ontario, L5L 1C6, Canada (Fei Luo).

## Abstract

The Oil Refinery (OR) consortium is a model methanogenic enrichment culture used to study anaerobic benzene degradation. Over half of the culture’s bacterial community consists of two closely related *Desulfobacterota* strains, designated ORM2a and ORM2b, whose mechanisms of benzene activation are unknown. Two new metagenomes, including a complete circularized metagenome-assembled genome (MAG) for ORM2a, enabled a thorough investigation of this culture’s proteome. Among the proteins identified were Bam-like subunits of an ATP-independent benzoyl-CoA degradation pathway, as well as downstream β-oxidation proteins yielding acetate. The most abundant proteins identified mapped to two ORM2a gene clusters of unknown function. Homologous and syntenic gene clusters were identified in genomes of ORM2b and a sulfate-reducing *Pelotomaculum* that also degrades benzene, as well as in nine contigs assembled from hydrothermal vent metagenomes. Extensive homology and structural predictions suggest that the first cluster – termed the “Magic” gene cluster – encodes for enzymes catalyzing the chemically challenging activation of benzene and subsequent transformation steps yielding benzoyl-CoA. The second (“Nanopod”) gene cluster encodes a transmembrane complex that may facilitate benzene transport across the cell membrane. Phylogenomic analyses place ORM2a and ORM2b within a novel genus of strict anaerobes specialized for benzene degradation, which we propose naming “*Candidatus* Anaerobenzenivorax”.

**IMPORTANCE:** Benzene is a widespread, persistent and toxic pollutant that can accumulate in anoxic environments such as groundwater and sediments. Despite decades of study, the biochemical mechanisms by which benzene is activated under anaerobic conditions remain unproven. This study provides strong genetic and proteomic evidence for a new class of enzymes that initiate anaerobic benzene activation and proposes a preliminary model for their underlying biochemistry. These findings lay a foundation for future biochemical studies and expand our understanding of how microbes carry out extreme redox chemistry in the absence of oxygen.

## INTRODUCTION

It has been 45 years since anaerobic benzene degradation was first documented by Ward et al. (1), reversing a long-held tenet that aromatic hydrocarbons were recalcitrant under anaerobic conditions. Notorious for its carcinogenicity, stringent environmental regulations, and persistence in anoxic environments, benzene has been the subject of numerous studies seeking to characterize its metabolism in the absence of oxygen (2, 3). Laboratory enrichment culture studies have confirmed unequivocally that anaerobic benzene degradation can be coupled to iron reduction, nitrate reduction, sulfate reduction, and methanogenesis, catalyzed by a handful of specialized microbial clades (2, 4). But despite great efforts, the biochemical mechanism(s) by which benzene is activated in the absence of oxygen remain elusive.

Initial studies using ^13^C- or ^14^C-benzene and occasionally H ^18^O identified three possible activation reactions based on the detection of labelled metabolites: (i) anaerobic hydroxylation of benzene yielding phenol (5–12), (ii) carboxylation of benzene yielding benzoate (6, 8–10, 13, 14), and (iii) methylation of benzene yielding toluene (10). However, neither phenol nor benzoate are diagnostic of a single activation mechanism (15), and supporting evidence of benzene methylation is sparse in literature (16, 17). From 2003-2008, hydrogen and carbon isotope fractionation ratios were evaluated in active benzene-degrading enrichment cultures (18–20). The slope of dual isotope plots (Δδ ^2^H/Δδ ^13^C, a value defined as Λ) from nitrate-reducing cultures (Λ = 8 to 19, *n* = 5) was substantially lower that than for sulfate-reducing and methanogenic cultures surveyed (Λ = 28 to 31, *n* = 7), hinting that at least two distinct biochemical benzene activation mechanisms exist in nature (18).

By 2010, advances in DNA sequencing and analysis enabled meta’omic characterization of anaerobic benzene degraders for the first time (21–23). Abu Laban *et al*. (24) sequenced the metagenome of the iron-reducing enrichment culture BF and identified a protein with homology to a known phenylphosphate carboxylase (PpcA). The corresponding putative anaerobic benzene carboxylation gene (*abcA*) was located on a contig affiliated with *Clostridia*. Expression of the putative *abcA* and neighboring genes (*abcD*, benzoyl-CoA ligase [*bzlA*] and a *ubiX*-like gene) was induced with benzene or benzoate, but not with phenol, suggesting a possible role in benzene activation (24). Syntenic gene clusters have since been identified in the transcriptomes of several nitrate-reducing cultures (25–27), including one previously analyzed by isotopic fractionation (18, 19). Most nitrate- and iron-reducing benzene degraders identified to date have been affiliated with the order *Thermincolales* (25–29); a putative benzene-carboxylating, sulfate-reducing *Thermincolales* was also recently identified in a jet fuel-contaminated aquifer via metagenomics (30). Researchers are now attempting to verify benzene carboxylation using structural and biochemical assays, among other tools (31).

Evidence for benzene hydroxylation has only been shown for one microorganism, *Geobacter metallireducens* GS-15, via gene deletion. In 2013-2014, Zhang *et al.* (5, 23) identified a handful of genes that were upregulated during active phenol or benzene metabolism, and deletion of key genes (Gmet0231, Gmet0232, or *ppsA*) inhibited benzene degradation. The gene *ppsA* encodes for the alpha subunit of phenylphosphate synthase, which catalyzes the first step in anaerobic phenol metabolism (32, 33). Functional annotations suggest that the products of Gmet0231 and Gmet0232 encode oxidoreductases that could potentially catalyze the addition of water to benzene, forming phenol. Further investigation into the exact functions of these enzymes is needed. In 2024, Bullows *et al.* (17) identified a fumarate addition gene cluster consisting of a 3-hydroxybenzylsuccinate synthase and a methyltransferase in the genome of *Geotalea daltonii* strain FRC-32, a close relative of *Geobacter*. Expression of these genes – previously known only to catalyze activation of alkanes and alkyl-substituted aromatic rings (34) – was upregulated during active benzene metabolism but not with toluene. Addition of benzene to *G. daltonii* whole-cell lysates resulted in the near-stoichiometric formation of toluene, but curiously benzene was completely depleted before toluene was formed (17). Whether this gene cluster truly encodes for benzene methylation enzymes remains uncertain.

Homologs of known and predicted anaerobic aromatic-activating genes have not been detected in methanogenic benzene-degrading enrichment cultures (4, 21) nor in the well-characterized sulfate-reducing enrichment culture BPL (9, 35). Genes associated with aerobic metabolism (monooxygenases) are not found in the major taxa in these mixed cultures, although recent growth experiments found that aerobic benzene degraders may reside in these cultures at trace abundances (36). Additionally, these enrichments do not possess genes for the first step of anaerobic growth on benzoate (benzoyl-CoA ligase) nor exhibit growth on benzoate (21, 35). To date, metagenomic and proteomic surveys of representative cultures have identified the presence and expression of genes implicated in the ATP-independent metabolism of benzoyl-CoA and downstream β-oxidation intermediates (21, 35). Given this information and previous isotope fractionation data (18, 19) indicating that the mechanism of benzene activation catalyzed by methanogenic cultures is distinct from that in nitrate-reducing cultures – where *abcA* genes are present and highly expressed (25–27), and carboxylation is likely – we surmise that a novel hydrocarbon activation mechanism catalyzing anaerobic transformation of benzene to benzoyl-CoA must exist in our methanogenic culture.

In this study, we integrated results from three proteomic experiments on the methanogenic benzene-degrading OR consortium. The culture, maintained on benzene for over 25 years, harbors two closely related strains of benzene-degrading *Desulfobacterota* (ORM2a and ORM2b) consistently detected in high abundance (4, 21, 36–41). Although the relative abundance of each ORM2 strain varies by subculture (39), together they account for 48-92% of bacteria in this consortium (Table S1). They are the only microorganisms whose growth directly coincides with anaerobic benzene degradation activity, and no substrate other than benzene has been shown to support their growth (4, 21, 36, 39). The OR culture also contains methanogenic archaea (*Methanothrix* and *Methanoregula*) and over 100 low abundance microorganisms with various predicted functions (39, 40, 42). Initial proteomic studies were conducted on the OR consortium in the early 2010’s (21) but results were difficult to interpret due to the lack of a sound metagenome and low protein yields. In the years since, improvements in culture maintenance (36, 39) and new sequencing efforts (40) have enabled the reconstruction of two high quality metagenomes and 74 metagenome-assembled genomes (MAGs; Table S2). The MAG of ORM2a was successfully closed (NCBI accession no. CP113000.1), providing a valuable blueprint for deciphering anaerobic benzene degradation. Here, we discovered that the most abundant proteins expressed by the culture mapped to a unique cluster of syntenic genes found only in other benzene-degrading obligate anaerobes and in a handful of metagenomic contigs from deep hydrocarbon-producing hydrothermal vents.

## RESULTS

### Features of the ORM2a MAG

The ORM2a MAG consists of a single circular chromosome of 3,300,546 bp with low GC skew and no discernible terminus region (Figures 1 and S1). General genome statistics are summarized in Table S3. To assess metabolic potential, coding sequences predicted by the NCBI Prokaryotic Genome Annotation Pipeline were uploaded to BlastKOALA for automatic KEGG Orthology (KO) assignment (43). Forty-six percent of genes identified (1,431 of 3,135) were assigned KO identifiers and mapped into metabolic pathways using the KEGG Mapper Reconstruct tool (44, 45). Key pathways and genes of interest are highlighted below and in Figure 2; additional pathway and genomic features are detailed in Tables S4-S9 and Supplementary Text S1.

**Figure 1.**
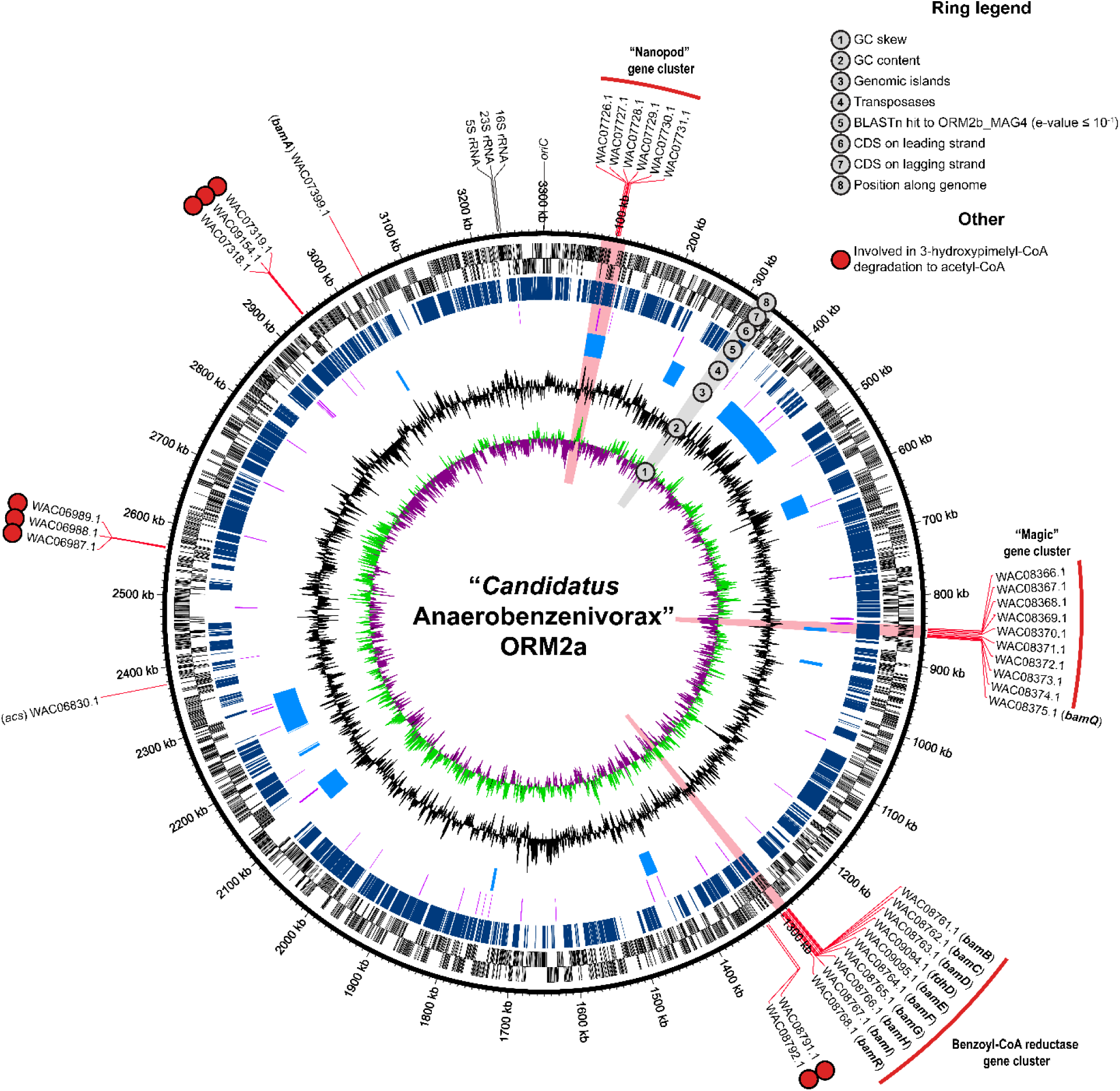
The complete ORM2a genome. Genomic neighborhoods and genes of interest, whose products were recovered with high certainty (≥99% confidence) from the OR proteomes, are highlighted in red and summarized in Table 2. Genes involved in the degradation of 3-hydroxypimelyl-CoA to acetyl-CoA are marked with red circles. Protein accession numbers are provided, and corresponding ORM2a gene locus tags are listed in Table S10. The positions of the origin of replication *(oriC*) and ribosomal RNA genes are also shown in this figure.

**Figure 2.**
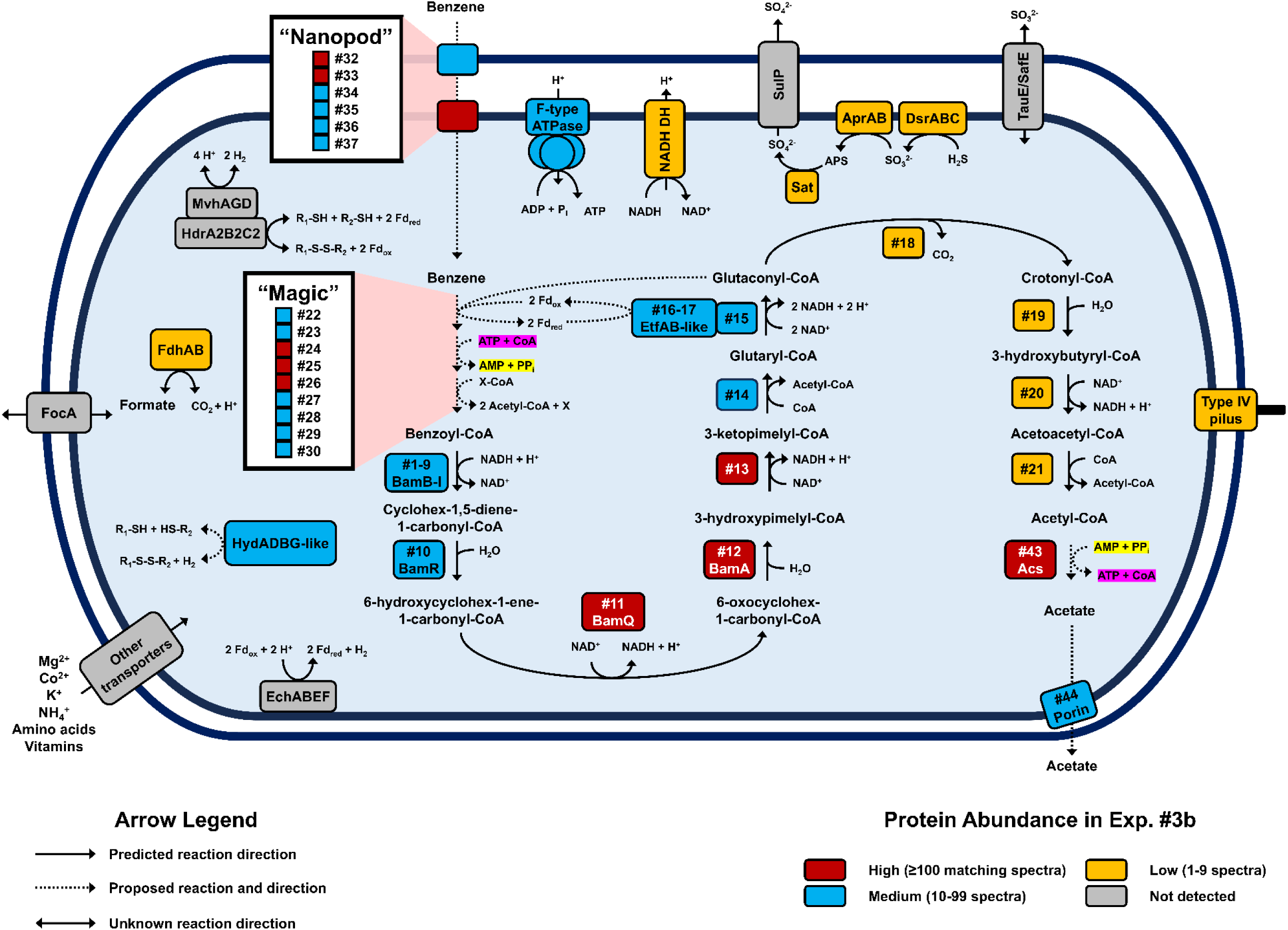
Summary of key metabolic pathways identified in the ORM2a proteome. Protein spectral counts from Experiment #3b were used to estimate protein abundances. Additional complete and incomplete metabolic pathways identified in the ORM2a genome are summarized in Table S4. Annotations for numbered proteins are provided in Table 2. Abbreviations (clockwise from top-center): ATPase, ATP synthase; NADH DH; NADH dehydrogenase; SulP, sulfate permease; Sat, sulfate adenylyltransferase; AprAB, adenylyl-sulfate reductase; Dsr, dissimilatory sulfite reductase; TauE/SafE, sulfite exporter; Acs, acetyl-CoA synthetase; EchABEF, energy converting hydrogenase; HydABGD-like, sulfhydrogenase; R-S-S-R, heterodisulfide; FocA, formate-nitrite transporter; FdhAB, formate dehydrogenase; Fd, ferredoxin; HdrA2B2C2, F_420_-reducing heterodisulfide reductase; Mvh, F_420_-non-reducing hydrogenase.

**Table 2.** Summary of high abundance proteins (≥100 matching spectra in Experiment #3b, bolded) and proteins of interest identified with ≥99% confidence. Proteins shown are mapped to one or more MAG reconstructed from OR metagenomes. Proteins are grouped by predicted biochemical functions and ranked by total spectra counts, with ties indicated by (=) symbols. Additional proteins identified are listed in Table S10.

| Protein # | Predicted function (NCBI Prokaryotic Genome Annotation Pipeline version 6.3 and best BLASTP hits) | Protein spectra count (Exp. #3b) | Protein abundance rank | Matching MAG(s) | Protein accession # |
| --- | --- | --- | --- | --- | --- |
| ***Benzoyl-CoA degradation to 3-hydroxypimelyl-CoA*** |  |  |  |  |  |
| 1 | Benzoyl-CoA reductase subunit BamB | 80 | =26 | ORM2a and ORM2b | WAC08761.1, ORM2b_MAG4_01005 |
| 2 | Benzoyl-CoA reductase subunit BamC | 15 | =154 | ORM2a and ORM2b | WAC08762.1, ORM2b_MAG4_01004 |
| 3 | Benzoyl-CoA reductase subunit BamD | 61 | =43 | ORM2a | WAC08763.1 |
| 4 | Formate dehydrogenase accessory sulfurtransferase FdhD | 6 | =287 | ORM2a | WAC09094.1 |
| 5 | Benzoyl-CoA reductase subunit BamE | 68 | 32 | ORM2a | WAC09095.1 |
| 6 | Benzoyl-CoA reductase subunit BamF | 18 | =135 | ORM2a | WAC08764.1 |
| 7 | Benzoyl-CoA reductase subunit BamG | 6 | =287 | ORM2a | WAC08765.1 |
| 8 | Benzoyl-CoA reductase subunit BamH | 91 | 23 | ORM2a and ORM2b | WAC08766.1, ORM2b_MAG4_04225 |
| 9 | Benzoyl-CoA reductase subunit BamI | 9 | =227 | ORM2a | WAC08767.1 |
| 10 | Cyclohexa-1,5-diene-1-carboxyl-CoA hydratase BamR | 49 | 59 | ORM2a | WAC08768.1 |
| 11 | **6-hydroxycyclohex-1-ene-1-carboxyl-CoA dehydrogenase BamQ** | **103** | **20** | **ORM2a** | **WAC08375.1** |
| 12 | **6-oxo-cyclohex-1-ene-carboxyl-CoA hydrolase BamA** | **143** | **13** | **ORM2a** | **WAC07399.1** |
| ***3-hydroxypimelyl-CoA degradation to acetyl-CoA*** |  |  |  |  |  |
| 13 | **3-hydroxyacyl-CoA dehydrogenase** | **148** | **9** | **ORM2a** | **WAC08792.1** |
| 14 | Thiolase family protein | 69 | 31 | ORM2a | WAC08791.1 |
| 15 | Glutaryl-CoA dehydrogenase (non-decarboxylating) | 58 | =49 | ORM2a | WAC06987.1 |
| 16 | Electron transfer flavoprotein subunit beta/FixA family protein | 39 | =69 | ORM2a | WAC06988.1 |
| 17 | Electron transfer flavoprotein subunit alpha/FixB family protein | 63 | =39 | ORM2a | WAC06989.1 |
| 18 | Glutaconyl-CoA decarboxylase subunit alpha | 5 | =332 | ORM2a | WAC08228.1 |
| 19 | Enoyl-CoA hydratase-related protein | 3 | =458 | ORM2a | WAC07318.1 |
| 20 | 3-hydroxybutyryl-CoA dehydrogenase | 9 | =227 | ORM2a | WAC09154.1 |
| 21 | Acetyl-CoA C-acetyltransferase | 3 | =458 | ORM2a | WAC07319.1 |
| ***Highly expressed "Magic" gene cluster putatively involved in benzene activation*** |  |  |  |  |  |
| 22 | AMP-binding protein | 74 | 29 | ORM2a and ORM2b | WAC08366.1, ORM2b_MAG4_03434 |
| 23 | MaoC family dehydratase | 57 | 52 | ORM2a | WAC08367.1 |
| 24 | **FAD-binding oxidoreductase** | **172** | **3** | **ORM2a** | **WAC08368.1** |
| 25 | **(Fe-S)-binding protein** | **219** | **2** | **ORM2a** | **WAC08369.1** |
| 26 | **FAD-binding oxidoreductase** | **265** | **1** | **ORM2a** | **WAC08370.1** |
| 27 | Type III CoA transferase | 45 | 61 | ORM2a | WAC08371.1 |
| 28 | Type III CoA transferase | 58 | =49 | ORM2a and ORM2b | WAC08372.1, ORM2b_MAG4_03441 |
| 29 | Hypothetical protein | 19 | =128 | ORM2a and ORM2b | WAC08373.1, ORM2b_MAG4_03442 |
| 30 | MaoC family dehydratase | 50 | =57 | ORM2a | WAC08374.1 |
| 31 | Transcription termination factor Rho | 25 | =100 | ORM2a | WAC08378.1 |
| ***Highly expressed "Nanopod" gene cluster putatively involved in benzene transportation across cell membrane*** |  |  |  |  |  |
| 32 | Hypothetical protein | 106 | =18 | ORM2a | WAC07726.1 |
| 33 | **DUF1329 domain-containing protein** | **158** | **4** | **ORM2a** | **WAC07727.1, ORM2b_MAG4_01802** |
| 34 | Universal stress protein | 6 | =287 | ORM2a | WAC07728.1 |
| 35 | Hypothetical protein | 7 | =269 | ORM2a | WAC07729.1 |
| 36 | YCF48-related protein | 65 | =36 | ORM2a | WAC07730.1 |
| 37 | MMPL family transporter | 60 | =45 | ORM2a | WAC07731.1 |
| ***Archaeal proteins associated with methanogenesis*** |  |  |  |  |  |
| 38 | Acetate-CoA synthetase Acs | 142 | 14 | *Methanotrix* MAG20 | MAG20_01236 |
| 39 | Coenzyme-B sulfoethylthiotransferase McrB | 149 | 8 | *Methanotrix* MAG21 | MAG21_01157 |
| 40 | Coenzyme-B sulfoethylthiotransferase McrB | 154 | 7 | *Methanotrix* MAG11, MAG20 and MAG28 | MAG11_02361, MAG20_02648, MAG28_00841 |
| 41 | Acetate-CoA synthetase Acs | 125 | 16 | *Methanotrix* | MAG11_00500 |
| ***Other proteins*** |  |  |  |  |  |
| 42 | Chaperonin GroL | 156 | =5 | ORM2a | WAC08671.1 |
| 43 | Acetate-CoA synthetase Acs | 156 | =5 | ORM2a | WAC06830.1 |
| 44 | Porin | 53 | 53 | ORM2a | WAC06831.1 |

The ORM2a MAG contains few and mostly incomplete pathways for metabolizing carbohydrates, lipids, amino acids, and nucleic acids. Similarly, pathways for electron transport and ATP generation are sparse, comprising an F-type ATPase, a Nuo-type NADH dehydrogenase with one missing subunit (*nuoG*), and a complete dissimilatory sulfate reduction pathway (Figure 2). This finding supports recent experimental evidence that the OR consortium can couple benzene oxidation to sulfate reduction in addition to methanogenesis (4). BLAST searches and manual inspections confirmed the absence of homologs to known hydrocarbon-activating genes and to benzoyl-CoA ligase. The latter is a significant distinction between microorganisms containing putative anaerobic benzene carboxylase genes like *abcA* and those that do not (35). The presence of a complete ATP-independent benzoyl-CoA degradation pathway is consistent with ORM2a’s involvement in anaerobic benzene oxidation (Figures 1 and 2).

Additional genomic evidence suggests that ORM2a is capable of fermentative metabolism in syntrophic association with methanogenic archaea (Figure 2). Three gene clusters encoding energy-converting hydrogenase (Ech) and formate dehydrogenase (Fdh) complexes were identified that may facilitate interspecies electron transfer (46, 47). The MAG also encodes for a cytoplasmic Mvh-type F_420_-non-reducing hydrogenase – potentially used for hydrogenotrophic growth on carbon dioxide (48, 49) – and a putative sulfur-reducing hydrogenase operon (Table S6). Sulfhydrogenases (also called sulfur-reducing hydrogenases) are poorly characterized enzymes that can catalyze the reduction of elemental sulfur (S⁰) or polysulfides (S_n_^2-^) to hydrogen sulfide (H_2_S) using H_2_ as an electron donor (50) (Figure 2). This process has been most thoroughly described in *Pyrococcus furiosus* and related thermophilic sulfur-oxidizing archaea (50–53). Under certain conditions, some sulfhydrogenases may also catalyze the reverse reaction, producing H_2_ (50, 52). In the OR culture, the major sulfur source in the growth medium is ferrous sulfide (FeS) used as a reductant (38). It is unknown whether sulfhydrogenases can oxidize FeS or reduce oxidized forms such as FeS_2_. Notably, no visible S^0^ or polysulfide formation (e.g., yellow solids) have ever been observed in the 25+ years this culture has been maintained.

A draft MAG belonging to ORM2b (designated ORM2b_MAG4) was assembled into 378 contigs and is estimated to be 74.6% complete (40). Although the exact taxonomic identity of ORM2b_MAG4 remains uncertain due to the absence of a 16S rRNA gene – precluding direct comparison to a reference clone sequence of ORM2b (NCBI accession no. KT025834) – the high average nucleotide identity (89.94%) between ORM2a and ORM2b_MAG4 supports its correct classification (Figure 1, ring 5; Figure S2). No other MAGs recovered from the OR consortium metagenomes display significant nucleotide identity to ORM2a.

### Proteomic analyses

Between 2010 and 2018, three proteomics experiments were performed on subcultures from the same OR culture lineage (OR-b1C, OR-b, and OR-b1A; see Figure S3). Experiments #1 (2010) and #2 (2011) used LC-MS/MS to analyze trypsin-digested crude extracts and recovered 153 unique proteins (Table 1) – a very low yield. To improve detection, protein separation by gel electrophoresis was introduced in Experiment #3 (2018). Duplicate crude extracts were prepared: one was directly digested with trypsin (Experiment #3a), as in previous protocols, and yielded similarly few proteins (Table 1). The second crude extract (Experiment #3b) was separated on a 15% SDS-PAGE gel, sliced into five equal sections (Figure S4), and subjected to in-gel trypsin digestion and LC-MS/MS. This method yielded over 800 unique proteins (Table 1).

**Table 1.**
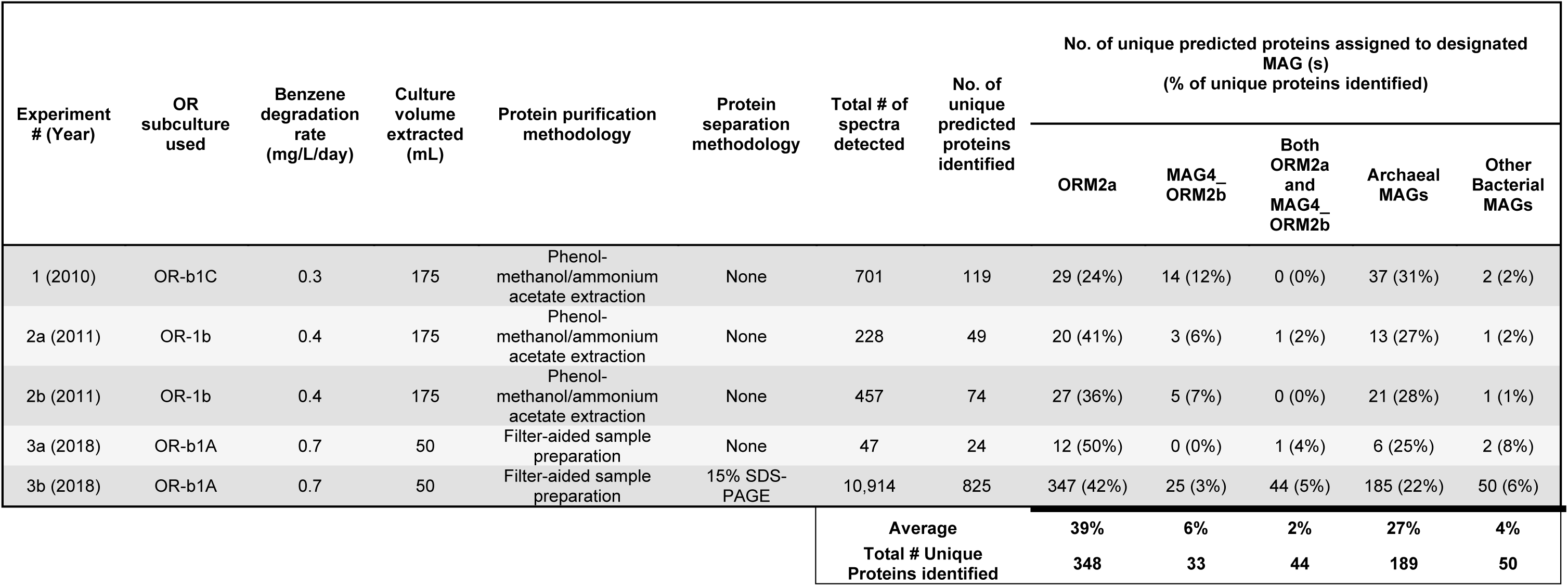
Summary of proteomic experiments performed on the Oil Refinery (OR) enrichment culture. Experiments differed in sample preparation methods, fractionation approaches, and instrumentation. Experiment #1 analyzed one protein sample, while Experiments #2 and #3 included technical replicates (a and b). All samples were subjected to protein extraction by sonication.

| Experiment # (Year) | OR subculture used | Benzene degradation rate (mg/L/day) | Culture volume extracted (mL) | Protein purification methodology | Protein separation methodology | Total # of spectra detected | No. of unique predicted proteins identified | No. of unique predicted proteins assigned to designated MAG (s)<br>(% of unique proteins identified) |  |  |  |  |
| --- | --- | --- | --- | --- | --- | --- | --- | --- | --- | --- | --- | --- |
|  |  |  |  |  |  |  |  | ORM2a | MAG4_ORM2b | Both ORM2a and MAG4_ORM2b | Archaeal MAGs | Other Bacterial MAGs |
| 1 (2010) | OR-b1C | 0.3 | 175 | Phenol-methanol/ammonium acetate extraction | None | 701 | 119 | 29 (24%) | 14 (12%) | 0 (0%) | 37 (31%) | 2 (2%) |
| 2a (2011) | OR-1b | 0.4 | 175 | Phenol-methanol/ammonium acetate extraction | None | 228 | 49 | 20 (41%) | 3 (6%) | 1 (2%) | 13 (27%) | 1 (2%) |
| 2b (2011) | OR-1b | 0.4 | 175 | Phenol-methanol/ammonium acetate extraction | None | 457 | 74 | 27 (36%) | 5 (7%) | 0 (0%) | 21 (28%) | 1 (1%) |
| 3a (2018) | OR-b1A | 0.7 | 50 | Filter-aided sample preparation | None | 47 | 24 | 12 (50%) | 0 (0%) | 1 (4%) | 6 (25%) | 2 (8%) |
| 3b (2018) | OR-b1A | 0.7 | 50 | Filter-aided sample preparation | 15% SDS-PAGE | 10,914 | 825 | 347 (42%) | 25 (3%) | 44 (5%) | 185 (22%) | 50 (6%) |
| **Average** |  |  |  |  |  |  |  | **39%** | **6%** | **2%** | **27%** | **4%** |
| **Total # Unique Proteins identified** |  |  |  |  |  |  |  | **348** | **33** | **44** | **189** | **50** |

In total, 858 unique proteins were identified across all experiments (Table 1). Protein accession numbers, automated annotations, peptide and spectral counts, and amino acid sequences are provided in Table S10, with summary statistics in Tables S11a and S11b. Proteins that mapped to one or more MAG, are putatively involved in anaerobic benzene degradation, and/or were detected in high abundance in Experiment #3b (≥100 spectra) are highlighted in Table 2 and Figures 1-4. Notably, the most high-abundant proteins detected in Experiment #3b were also observed in Experiments #1 and #2 (Table S10), demonstrating strong reproducibility of our results despite years between tests and differences in extraction and analysis methods (Table 1).

### Key microorganisms identified in the OR proteomes

Half of all proteins identified mapped to the MAGs of ORM2a and/or ORM2b (Tables 1 and 2). Of these, 348 proteins mapped exclusively to ORM2a (41%), 33 proteins mapped exclusively to ORM2b_MAG4 (4%), and 44 proteins (5%) mapped to both MAGs (Table 1). Most ORM2b_MAG4 proteins identified shared 71-100% amino acid identity to a homologous ORM2a protein or undetected ORM2a gene products. Only six low abundance proteins – associated with protein synthesis or hypothetical functions – were unique to ORM2b_MAG4 (Table S10). Given these similarities, the following sections focus on ORM2a proteins and metabolic pathways of interest.

One-quarter of proteins identified mapped to MAGs of methanogenic archaea (Tables 1 and S10), specifically acetoclastic *Methanothrix* spp. (170 proteins across 8 MAGs) and hydrogenotrophic *Methanoregula* spp. (19 proteins across 3 MAGs; ∼89% fewer than *Methanothrix*). Most high abundance archaeal proteins were affiliated with methanogenesis and mapped to more than one MAG (Table 2). Forty-four low-abundance proteins (5%) mapped to other bacterial MAGs including “*Ca.* Nealsonbacteria” DGGOD1a. The remaining proteins could not be assigned to a specific MAG (180 proteins, 21%) or mapped to contaminant/decoy sequences (20 proteins, 2%).

### Benzoyl-CoA degradation to acetyl-CoA

We next searched for enzymes involved in downstream aromatic degradation pathways and detected a complete suite of putative benzoyl-CoA degradation proteins encoded on the ORM2a genome (Figures 1 and 2; Table 2). Ten gene products (designated Proteins #1-10) mapped to a single gene cluster and were annotated as subunits of a Bam-type benzoyl-CoA reductase (BamBCDEFDHI; whose components are described in Wischgoll et al. (54) and Boll et al. (55)), an FdhD-like insertase, and a cyclohexa-1,5-diene-1-carbonyl-CoA hydratase (BamR). Two high abundance proteins annotated as 6-hydroxycyclohex-1-ene-1-carbonyl-CoA dehydrogenase (BamQ, Protein #11) and 6-oxo-cyclohex-1-ene-carbonyl-CoA hydrolase (BamA, Protein #12) were encoded elsewhere in the ORM2a genome (Figure 1). Multiple sequence alignments (Table S12) and maximum likelihood phylogenetic tree analyses (Figure S5) support the functional annotation of all predicted Bam proteins. The only other proteins identified that likely participate in anaerobic benzoyl-CoA degradation were three *bam* genes mapping to ORM2b_MAG4 (Table 2). These results corroborate longstanding evidence that ORM2a and ORM2b are key benzene-degrading members of the OR consortium (4, 21, 36, 39, 41).

Nine proteins predicted to catalyze the transformation of 3-hydroxypimelyl-CoA to acetyl-CoA were also identified and mapped exclusively to the ORM2a MAG (Figures 1-2; Table 2). In our most successful proteome (Experiment #3b), proteins involved in converting 3-hydroxypimelyl-CoA to glutaconyl-CoA (Proteins #13-17) had higher spectral counts than downstream beta-oxidation enzymes (Proteins #18-21). This observation may indicate a metabolic branch point (Figure 2), as discussed later. Notably, the gene encoding a non-decarboxylating glutaryl-CoA dehydrogenase (Protein #15) is located in the same operon as two genes encoding bifurcating electron transfer flavoproteins (EtfAB-like; Proteins #16-17) (Table 2). EtfAB typically mediate flavin-based electron bifurcation and confurcation reactions, which are used to drive an endergonic redox reaction by coupling it with an exergonic one (56). As illustrated in Figure 2, the EtfAB-like complex may couple the endergonic reduction of NAD^+^ with the exergonic oxidation of glutaryl-CoA and ferredoxin, in a manner analogous to EtfAB-butyryl-CoA dehydrogenase (56, 57).

### Sulfur metabolism and acetate production

Several unexpected ORM2a proteins were detected in the OR proteomes, including six associated with the dissimilatory reduction of sulfate to hydrogen sulfide – specifically sulfate adenylyltransferase (Sat), adenylylsulfate reductase (AprAB), and dissimilatory sulfite reductase components (DsrAB, DsrC) – despite not adding any sulfate to the culture medium (Figure 2). Additionally, products of the putative sulfhydrogenase operon (HydADBG) were the only hydrogenase proteins detected that mapped to the ORM2a genome (Figure 2 and Table S6). If this enzyme consumes H_2_ released during benzene oxidation, it could explain why so few proteins from hydrogenotrophic methanogens (*Methanoregula* spp.) were recovered from the OR proteomes (Table S10). Together, these findings raise new questions about ORM2a’s sulfur metabolism that warrant further investigation.

The protein folding chaperonin GroEL (Protein #42) and a putative AMP-forming acetyl-CoA synthetase (Acs, #43) were tied as the 5^th^ most abundant proteins in Experiment #3b (Table 2). Acs normally catalyzes the ATP-dependent formation of acetyl-CoA molecules from acetate and coenzyme A but given that acetyl-CoA is already produced in abundance during anaerobic benzene metabolism, we surmise that Protein #43 catalyzes the reverse reaction yielding acetate (a substrate for acetoclastic *Methanothrix* spp.), coenzyme A, and ATP (Figure 2). Similar reverse reactions were proposed for the anaerobic benzoate degrader *Syntrophus acidotrophicus* (58, 59) and three syntrophic bacterial MAGs reconstructed from full-scale anaerobic digesters (60). None of these genomes code for acetate kinase or closely related homologs needed to phosphorylate acetate. ORM2a also lacks cytosolic pyrophosphatase but harbors a membrane-bound Na⁺-translocating pyrophosphatase (WAC07798.1), suggesting similarly elevated intracellular pyrophosphate levels that may favor ATP production in syntrophic organisms (59). Phylogenetic trees support the correct annotation of Protein #43 (Figure S6). An unclassified porin (Protein #44) whose corresponding gene is localized adjacent to *acs* may facilitate acetate export from ORM2a cells (Figure 2).

### Highly expressed ORM2a proteins from gene clusters of unknown function

The three most abundant and consistently detected proteins in all OR proteomes (Proteins #24-26) mapped consecutively to an ORM2a gene cluster of unknown function (Table 2; Figures 1, 3 and 4a). Products of several neighboring genes (Proteins #22-23, #27-30) were also detected in multiple OR proteomes. This region, designated herein as the “Magic” gene cluster, is located on a small genomic island flanked by multiple predicted transposases and pseudogenes (Figure 1; Table S7). Five operons were predicted within the “Magic” gene cluster, each encoding for two to three proteins (Figure 4a). Automated annotations from NCBI and IMG provided few insights into their functions (Table 2) with one notable exception: the tenth consecutive gene in this cluster most likely encodes 6-hydroxycyclohex-1-ene-1-carbonyl-CoA dehydrogenase (BamQ, Protein #11), a known component of the benzoyl-CoA degradation pathway (Figure 2). The detection of a neighboring transcription-terminating Rho factor in Experiment #3b (Protein #31) suggests that expression of the “Magic” gene cluster is tightly regulated.

**Figure 3.**
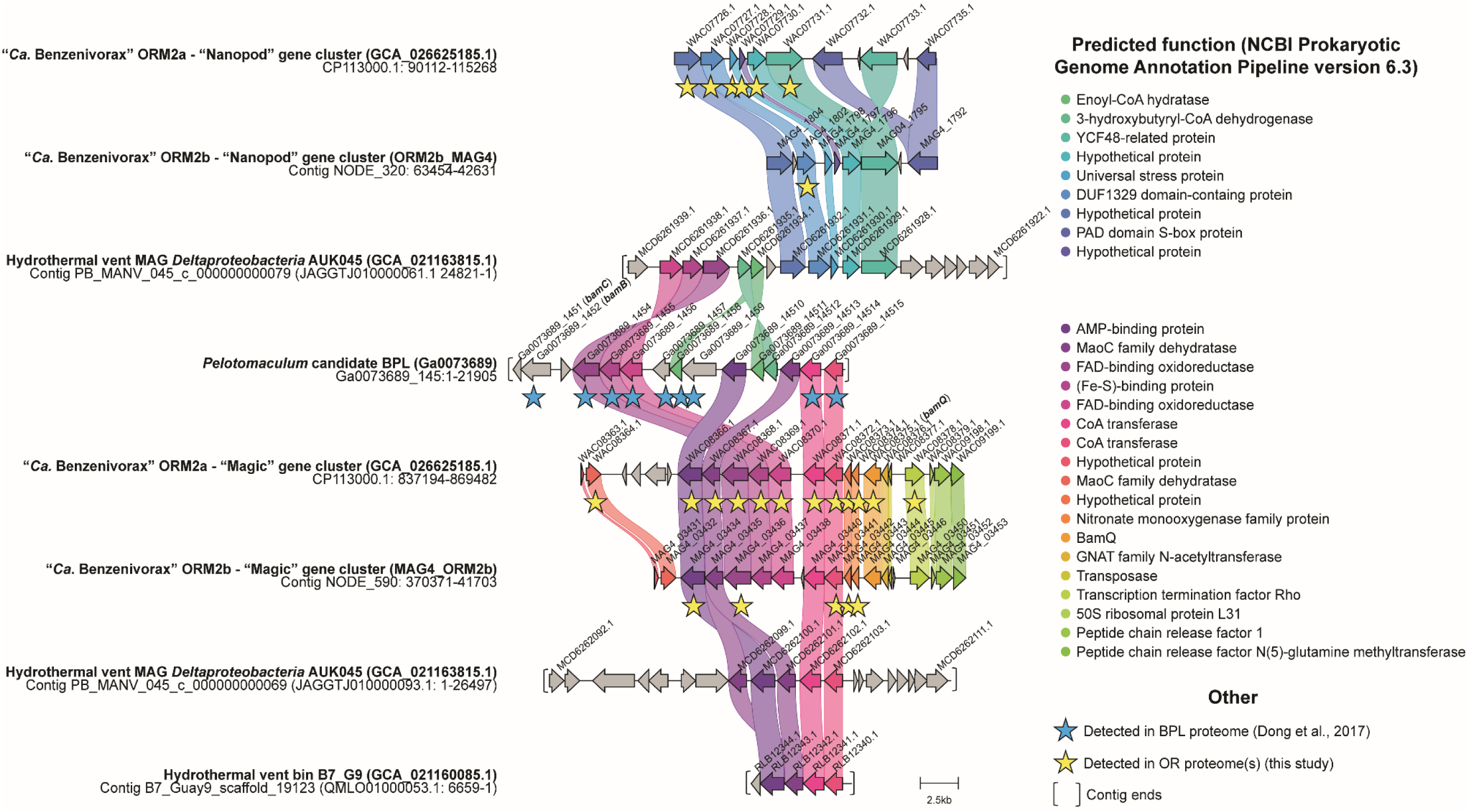
Genomic neighborhoods of all publicly available MAGs in NCBI and IMG producing significant alignments to the “Magic” and “Nanopod” gene clusters in ORM2a and ORM2b_MAG4. Homologous genes (≥40% amino acid identity) are connected and color-coded by function; sequence alignment statistics for relevant proteins are shown in Table 3 and additional homology statistics are provided in Table S12. Corresponding products detected in the proteomes of the OR consortium and the enrichment culture BPL are marked with yellow or blue stars, respectively. The ends of a contig are capped with brackets.

**Figure 4.**
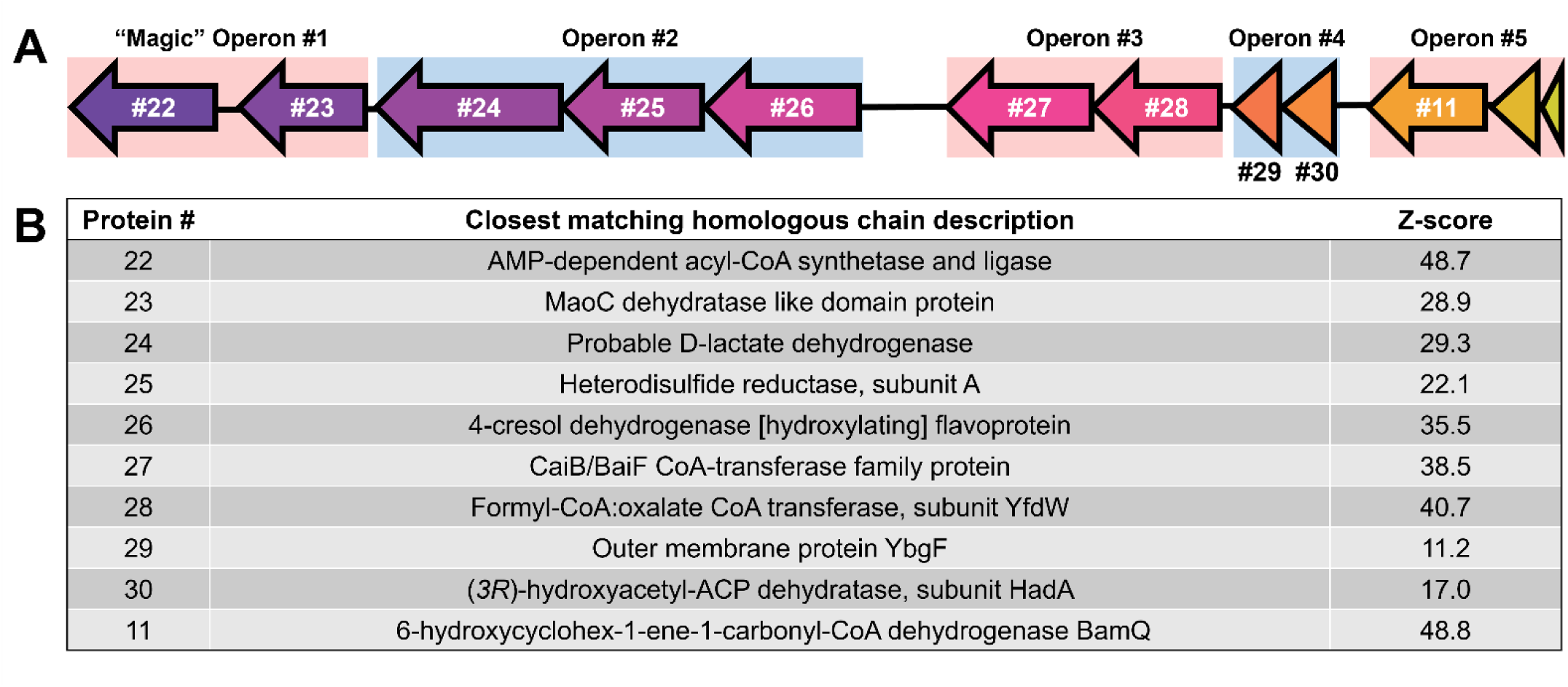
Structure and functional prediction of proteins encoded by the “Magic” gene cluster. Panel A shows the operon organization of this gene cluster. Panel B summarizes the best Protein Data Bank (PDB) search results for each “Magic” gene product. Z-scores are a measure of protein composite confidence, and Z-scores ≥30 are considered significant. More alignment statistics are provided in Table S13.

**Table 3.** BLASTP search results for ORM2a “Magic” and “Nanopod” proteins against MAG4_ORM2b and publicly available sequence databases in GenBank and JGI (as of August, 2024). NCBI accession numbers and percent amino acid identities are shown for best BLASTP hits to select MAGs and bins with significant alignments (e-value < 1.0E-05, coverage ≥ 50%) to ORM2a proteins. Protein accession numbers and best BLASTP hit search results for other ORM2a proteins of interest and other reference genomes are provided in Table S12.

| Protein cluster # | ORM2a | ORM2b_MAG4 | *Pelotomaculum* candidate BPL MAG | Hydrothermal vent MAG AUK045 | Hydrothermal vent bin B94_G9 | Hydrothermal vent bin B7_G9 | Hydrothermal vent MAG, Pescadero Basin | Wastewater MAG, Germany | Freshwater sediment MAG, Oklahoma |
| --- | --- | --- | --- | --- | --- | --- | --- | --- | --- |
|  | This study | This study | Ga0073689 (IMG) | GCA_021163815.1 | GCA_003646755.1 | GCA_003646785.1 | GCA_021160085.1 | GCA_017996625.1 | GCA_016935335.1 |
| *Highly expressed "Magic" gene cluster putatively involved in benzene activation* |  |  |  |  |  |  |  |  |  |
| 23 | 100% | 96% | 76% | 72% | 71% | 71% | 25% | 24% | 27% |
| 24 | 100% | 95% | 76% | 73% |  | 73% |  |  |  |
| 25 | 100% | 97% | 74% | 76% |  |  |  | 22% |  |
| 26 | 100% | 97% | 70% | 69% | 66% | 28% | 26% | 29% | 29% |
| 27 | 100% | 94% | 66% | 62% | 61% |  |  | 24% |  |
| 28 | 100% | 98% | 74% | 65% | 72% | 69% |  |  | 28% |
| 29 | 100% | 98% | 69% | 60% | 62% | 62% |  |  | 29% |
| 30 | 100% | 90% |  | 61% | 51% |  |  |  |  |
| 31 | 100% | 90% |  | 60% | 52% |  |  |  |  |
| 11 | 100% | 94% | 64% | 27% |  | 57% | 21% | 62% |  |
| *Highly expressed "Nanopod" gene cluster putatively involved in benzene transportation across cell membrane* |  |  |  |  |  |  |  |  |  |
| 32 | 100% | 84% |  | 44% |  | 26% | 30% | 23% | 29% |
| 33 | 100% | 94% |  | 60% | 32% | 36% | 31% | 23% | 33% |
| 34 | 100% | 82% | 32% | 43% | 31% | 29% |  | 28% | 31% |
| 35 | 100% | 92% |  |  |  |  |  |  |  |
| 36 | 100% | 88% |  | 46% |  |  | 31% | 30% | 30% |
| 37 | 100% | 92% |  | 51% | 33% |  | 35% | 36% | 34% |

Two other highly abundant proteins (#32 and #33, ranked 18^th^ and 4^th^ in Experiment #3b, respectively) mapped to a second ORM2a genomic island of unknown function, designated the “Nanopod” gene cluster (Tables 2 and S7; Figures 1-3), which includes four other genes whose products were detected. Nanopods are tubular structures (<100 nm in diameter) that may facilitate nutrient transport across cell membranes or export toxicants (61, 62). Shetty et al. (61) found that phenanthrene induced nanopod formation in *Delftia acidovorans* Cs1-4 but not structurally similar hydrocarbons (naphthalene and toluene), and that deletion of its DUF1329 gene (DelCs14_1722) impaired growth on phenanthrene. BLASTP alignments suggest that Protein #33 is distantly related to the translated DelCs14_1722 gene sequence (21% amino acid identity), but the high expression of this ORM2a gene cluster and its possible relation to nanopod formation is intriguing.

### Comparative analyses of the “Magic” and “Nanopod” gene clusters to other genomes and proteomes

To determine whether other organisms harbor syntenic regions to the “Magic” and “Nanopod” gene clusters, translated protein sequences were searched against all reference genomes in NCBI and IMG databases (as of August 2024) using BLASTP. Four syntenic contigs were identified (Figure 3; Table 3), along with six shorter contigs (comprising 2-3 genes) containing homologous genes with 51-76% amino acid identity to those in the “Magic” gene cluster (Table S12). These genes also appear to be grouped into operons identical to those in the ORM2a genome (Figures 3-4a). Nine of the ten contigs identified were affiliated to *Deltaproteobacteria* spp. from hydrothermal vent systems (63, 64). Hydrocarbons are important energy sources for microbial communities in these ecosystems and are produced primarily by geological processes (65). Hydrothermal vent MAG *Deltaproteobacteria* AUK045 is notable for containing syntenic contigs to both the “Magic” and “Nanopod” gene clusters (Figure 3; Table 3). Additionally, genes encoding predicted downstream benzoyl-CoA degradation and beta-oxidation pathway components were identified in AUK045 and two other draft *Deltaproteobacteria* bins (Table S12), suggesting metabolic potential for aromatic compound degradation.

*Pelotomaculum* candidate BPL (IMG GOLD accession number Ga0073689), the dominant benzene degrader in a sulfate-reducing enrichment culture from Poland (9, 35), provides another compelling comparison. Dong et al. (35) reconstructed this genome in 2017 from metagenome data (2,986,7007 bp across 97 contigs, ∼99% completeness) and published concurrent proteomic data from an active benzene-degrading subculture. We identified a highly expressed *Pelotomaculum* contig (Ga0073689_145) producing eight significant alignments to the “Magic” gene cluster, exhibiting 64-76% amino acid identity and e-values ≤ 1.0E-178 (Figure 3; Table 3). The Ga0073689_145 contig also encodes a predicted benzoyl-CoA reductase subunit (BamB) and an enoyl-CoA hydratase, drawing parallels to the “Magic” gene neighborhoods in the ORM2a, ORM2b, and *Deltaproteobacteria* AUK045 MAGs (Figure 3). These genomic and proteomic parallels are striking and suggest a conserved role for the “Magic” cluster in anaerobic benzene degradation.

### Predicted functions of the “Magic” proteins

Multiple sequence- and structure-based protein prediction tools were used to investigate the potential biochemical roles of proteins encoded by the “Magic” and “Nanopod” gene clusters. Search results for the “Magic” proteins are summarized below, in Figure 4b and in Tables S13-S14. Structure predictions for the “Magic” proteins and putative enzyme complexes are shown in Figures 5 and S7-S8. Phylogenetic trees were also constructed for each protein; however, most were too distantly related to characterized enzymes to yield meaningful functional insights.

**Figure 5.**
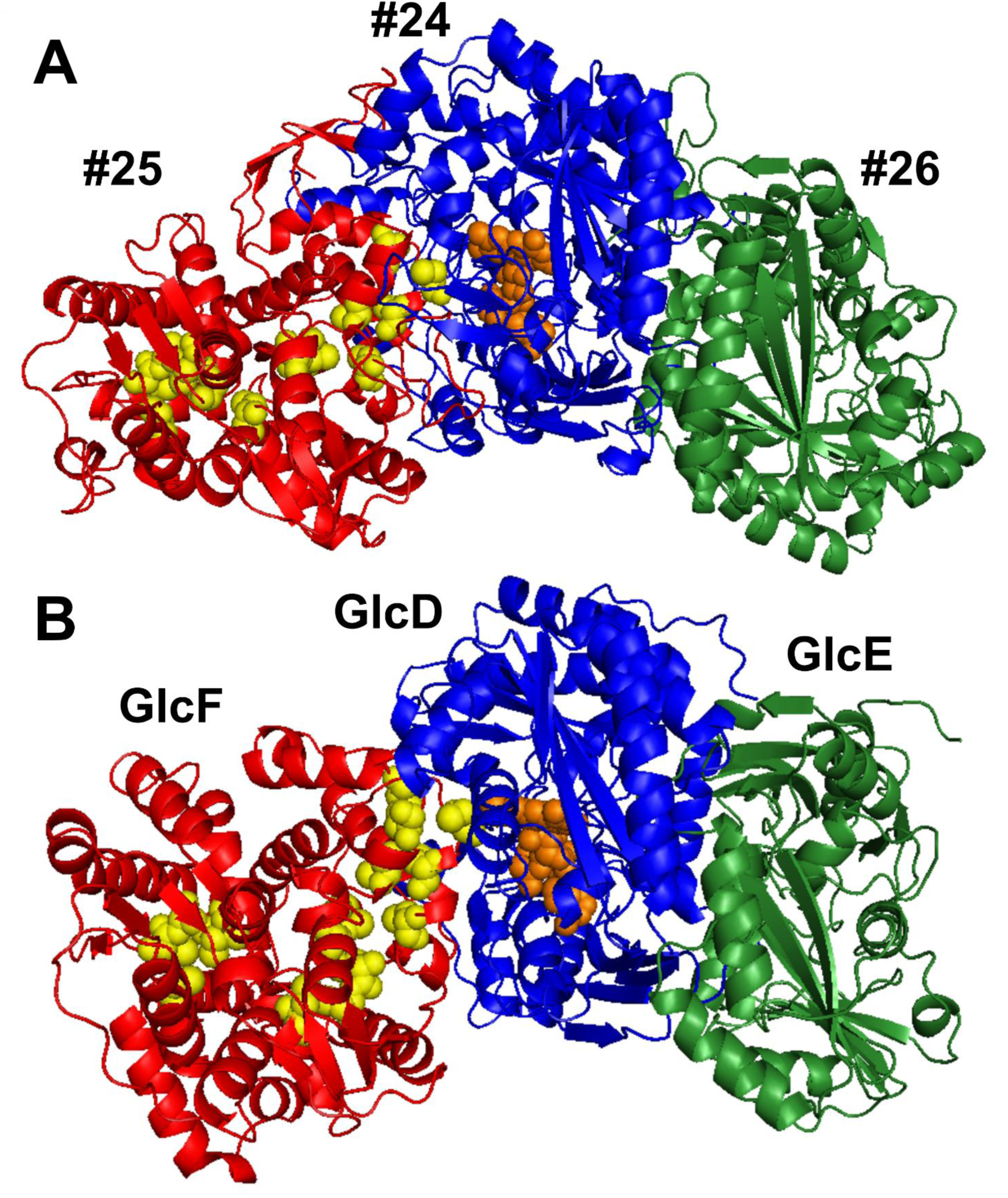
AlphaFold 3 predicted model of A) the Protein #24-26 enzyme complex and B) the glycolate oxidase complex GlcDEF in *Escherichia coli*. Homologous subunits are shown in red, blue, or green. FAD is rendered as orange spheres, and conserved cysteine residues potentially binding [4Fe–4S] clusters are shown as yellow spheres. A putative binding pocket is located near the FAD-binding site in Protein #24, which can be seen in Figure S8a. Structure confidence scores are shown in Figure S8b.

Operon #1 of the “Magic” gene cluster encodes Proteins #22 and #23 (Figure 4a). All prediction tools converged on the annotation of Protein #22 as a novel AMP-forming acyl-CoA synthetase (Figure 4b). Phylogenetic analysis placed Protein #22 and its homologs from other anaerobic benzene degraders in a distinct subfamily of acyl-CoA synthetases, separate from well-characterized enzymes such benzoyl-CoA ligase and acetyl-CoA synthetase (Figure S6). DeepFRI Gene Orthology (GO) predictions (66) further suggested that Protein #22 can bind to organic cyclic compounds such as benzene (Table S14b). Since Proteins #22 and #43 are predicted to use similar cofactors (e.g., ATP/AMP, CoA), these reactions may be coupled, with one reaction’s energy driving the other (Figure 2). Protein #23 contains a canonical “hot dog” domain typical of (*R*)-specific enoyl-CoA hydratases (67) and is likewise predicted to bind cyclic organic molecules (Table S13 and S14b). Enoyl-CoA hydratases catalyze the second step of β-oxidation (68), converting trans-2-enoyl-CoA molecules into 3-hydroxyacyl-CoA. Structure predictions for Proteins #22 and #23 are shown in Figures S7a and S7b.

Operon #3 encodes Proteins #27 and #28, predicted to form a class III CoA-transferase complex (Figures 4b and S7c). These enzymes operate via a ping-pong mechanism to transfer CoA groups between carboxylic acids. They are known to act on diverse substrates including oxalate and benzylsuccinate, which is the fumarate addition product of toluene (69). Proteins #29 and #30, encoded on Operon #4, were poorly annotated by most tools but may form a separate enzyme complex (Figures 4b and S7d). Interestingly, MotifFinder (70) detected weak homology in both proteins to domains in PaaZ, a bifunctional ring-hydrolyzing enzyme involved in phenylacetyl-CoA degradation (71) (Table S13). Operon #5 encodes BamQ (Figure 4b) and two proteins – a GNAT family N-acetyltransferase and a transposase-associated protein – that were not detected in the proteome (Figure 3). Homologs of these two proteins are also not encoded by the MAGs of *Pelotomaculum* candidate BPL or *Deltaproteobacteria* AUK045, suggesting they may not be involved in benzene degradation.

Proteins #24-26, encoded on Operon #2, were the three most abundant proteins detected in the OR proteomes (Figure 4a; Table 2). AlphaFold 3 predictions (72) suggest that these proteins form a heterotrimeric complex structurally similar to glycolate dehydrogenase in *Escherichia coli* (Figure 5). This enzyme complex is composed of two putative flavoproteins (GlcDE) and an Fe-S protein (GlcF), which together catalyze the oxidation of glycolate to glyoxylate (73). GlcD likely mediates substrate oxidation via its FAD cofactor and transfers electrons to an unknown acceptor through GlcF, a homolog of heterodisulfide reductase subunit HdrB (73, 74). HdrB reduces the coenzyme M-coenzyme B heterodisulfide to regenerate thiol cofactors, and is critical for methanogenesis (75). The electron acceptor(s) for bacterial HdrB homologs is unknown (76). The function of GlcE is also unknown, though knockout mutations to its corresponding gene abolish glycolate oxidation (73). Although the overall trimeric structures of GlcDEF and Proteins #24-26 are similar (Figure 5), their homologous subunits share ≤23% amino acid identity and likely catalyze distinct reactions.

Most prediction tools agreed that Proteins #24 and #26 are flavoproteins that may oxidize 2-hydroxy acids such as glycolate and D-lactate (Figure 4b). DeepFRI (66) predicted that both proteins can bind cyclic organic compounds, and that Protein #24 may also bind FAD (Table S13b) – a prediction supported by AlphaFold 3 models (Figure 5a). Intriguingly, a putative binding pocket is located adjacent to the FAD-binding site in Protein #24 (Figure S8). Phylogenetic analyses placed Protein #25 in an uncharacterized clade of Hdr-like enzymes (Figures S9 and S10). GO term predictions from PROST (77) suggest that it may have 2-hydroxyacid dehydrogenase and ferredoxin hydrogenase activity (Table S14a). Protein #25 contains four cysteine-rich motifs that could bind to electron-transferring Fe-S clusters (Figures 5a), akin to GlcF and HdrB (73, 74). However, two of the binding motifs lack key conserved cysteine residues, which might impair or prevent electron transfer to ferredoxin or other electron acceptors (Figure S11). Overall, functional inferences suggest that the “Magic” gene cluster encodes enzymes that act on aromatic structures, are rich in Fe-S clusters and FAD binding domains and likely convert benzene stepwise to benzoyl-CoA (see discussion).

### Predicted function of the “Nanopod” proteins

Protein prediction tools suggest that the “Nanopod” gene cluster encodes a transmembrane complex (Figure 2, Tables S13-S14). Protein #32 most closely resembles a porin, involved in passive substrate transport, whereas Protein #37 (a predicted RND superfamily efflux protein) appears to be involved in active transport. GO term searches in PROST (77) and DeepFRI (66) suggest that Protein #37 and possibly Protein #34 (an uncharacterized cytoplasmic protein) can bind to organic cyclic structures (Tables S14a and S14b). Most tools agreed that Protein #35 is a universal stress factor. Membrane stress can result from partitioning of hydrophobic compounds such as benzene into lipid bilayers (78). Collectively, it is plausible that the “Nanopod” gene cluster facilitates benzene movement across cell membranes, though the direction(s) of movement is not yet known (Figure 2).

### Phylogenomic placement of the ORM2a and ORM2b_MAG4 genomes

The genomes of ORM2a and ORM2b have undergone multiple reclassifications in response to ongoing taxonomic revisions (4, 39, 40). Our most recent phylogenomic analyses (Figure 6a) placed ORM2a and ORM2b within an uncharacterized clade of *Desulfobacterota*, distinct from previously assigned taxonomic groups including the candidate order Sva0485 and GTDB classes WTBG01 and UBA8473, as well a clade including *Deltaproteobacteria* MAG AUK045. The ORM2a and ORM2b clade – which appears to represent a novel class within the *Desulfobacterota* – also includes three environmental MAGs, one of which (GCA_021160085.1) was recovered from yet another hydrothermal vent system (63). While these MAGs harbor some downstream functional potential to degrade benzoyl-CoA and its intermediates (Table S12), they lack key genes implicated in benzene activation (Table 3).

**Figure 6.**
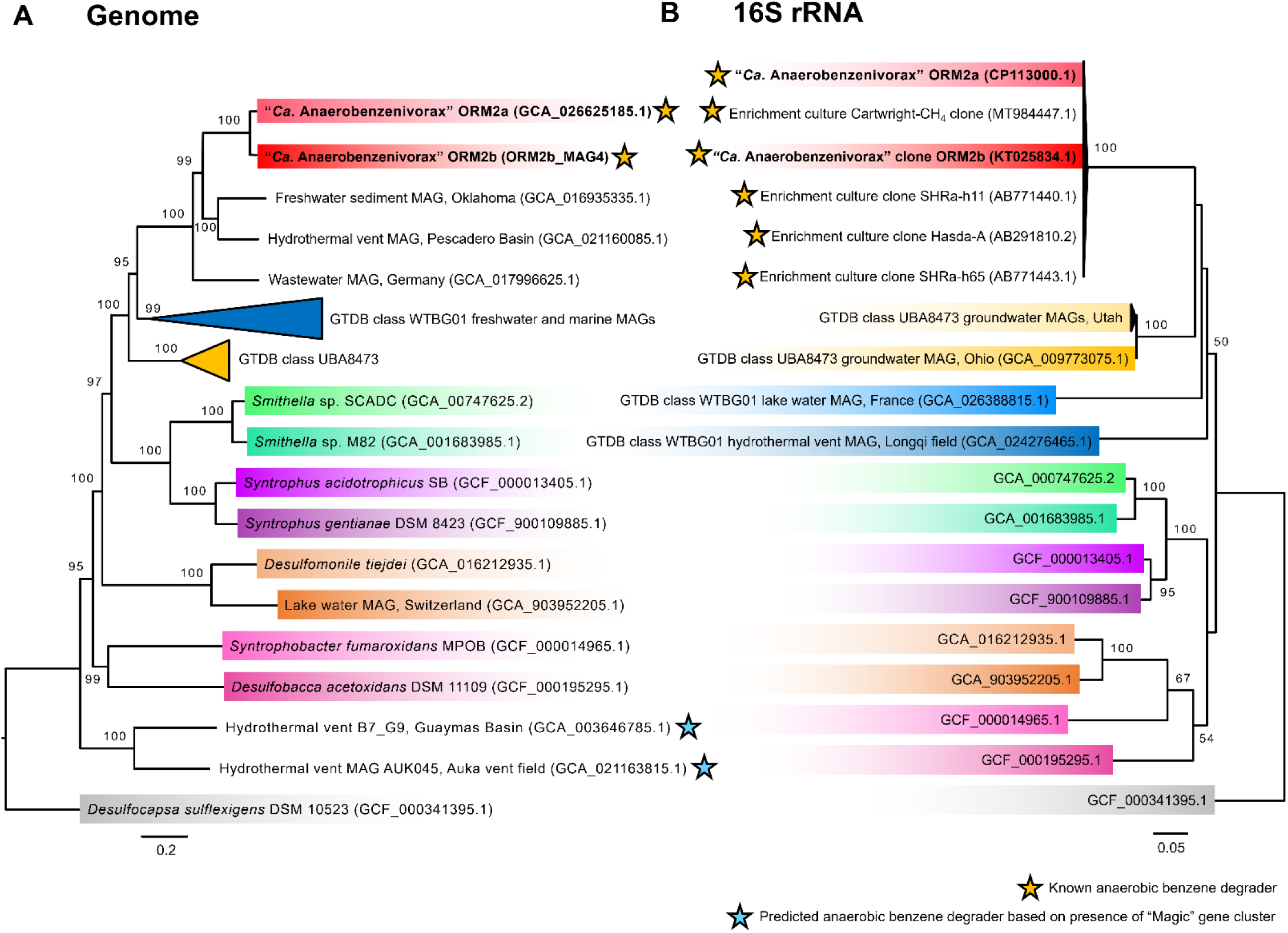
Taxonomic classification of ORM2a and reference *Desulfobacterota* species based on A) genome and B) 16S rRNA gene maximum likelihood phylogenetic trees. Sequences from this study are highlighted in bold. *Desulfocapsa sulflexigens* was used to root both trees. Bootstrap values <60% are not shown.

A complementary 16S rRNA gene-based phylogenetic tree (Figure 6b) revealed a consistent clustering pattern, clearly indicating that ORM2a and ORM2b form a monophyletic group of benzene-degrading specialists (4). These organisms – enriched from geographically diverse origins – share ≥ 99% 16S rRNA gene sequence identity and have only been observed to degrade benzene under sulfate-reducing and/or methanogenic conditions (38, 79, 80). Unfortunately, full genomes are lacking for most of these organisms. Taken together, the phylogenomic evidence presented here strongly supports the delineation of ORM2a and ORM2b as members of a novel genus within the *Desulfobacterota*. We propose the name “*Candidatus* Anaerobenzenivorax” (An.ae.ro.ben.ze.ni.i.vor’ax. Gr pref. *an*, not or without; Gr. n. *aer*, air; N.L. neut. n. *benzenum*, benzene, L. adj. *vorax,* devouring, N.L. neut. n. Anaerobenzenivorax, benzene-devouring without air). The proposed taxonomy is currently being registered in SeqCode (81).

## DISCUSSION

Many efforts have been made to elucidate mechanisms underlying anaerobic benzene activation. Recent advances in sequencing accuracy, protein detection, and *in silico* protein structure prediction are now enabling deeper insights into these processes. Because benzene is the only known growth-supporting substrate for organisms like “*Ca.* Anaerobenzenivorax” strains ORM2a and ORM2b, conventional differential growth experiments are not feasible. Instead, we must rely on proteomic snapshots during active benzene metabolism. As shown in this study, however, consistent detection of a highly expressed, conserved gene cluster – across multiple years and independent laboratories – strongly implicates this “Magic” gene cluster in initiating benzene metabolism via a novel mechanism that leads to benzoyl-CoA formation. Although biochemical validation is still required, we offer several early postulates regarding the core biochemistry of this elusive pathway.

### Proposed mechanism of anaerobic benzene activation catalyzed by ORM2a

The high activation energy required to functionalize benzene in the absence of oxygen is a long-standing challenge. Based on the enzymes identified herein and the example of toluene activation via fumarate addition (82, 83), we explored options for a radical mechanism. We propose that Proteins #24-26 form a novel oxidoreductase complex that catalyzes the initial activation of benzene via flavin-based radical chemistry. The radical flavin is most likely located in Protein #24 (Figure 5a) and transfers electrons to Protein #25, similar to GlcF and heterodisulfide reductase subunits (Figures S9–S10), but with mutations in two of its four cysteine-rich motifs that may preclude the use of an external molecule as an electron sink (Figure S11).

In our conceptual model (Figure 7), benzene is attacked by a radical generated by flavin-mediated single-electron oxidation of glutaconyl-CoA, a downstream intermediate of anaerobic benzene degradation. Flavin-mediated generation of an acyl-CoA radical was previously proposed for 4-hydroxybutyryl-CoA dehydratase (84). In ORM2a, proteomic evidence supports the idea that glutaconyl-CoA metabolism diverges into two distinct branches: one that proceeds through beta-oxidation reactions to yield acetyl-CoA (via Proteins #18-21), and another – proposed here – that uses glutaconyl-CoA as an acceptor in the initial benzene activation step (Figures 2 and 7). The addition reaction could occur across *para* positions of the aromatic ring, yielding a cyclohexadienyl-type radical intermediate. Resonance stabilization and internal rearrangement would produce 3-phenylglutaryl-CoA, with the radical localized to the α-carbon adjacent to the CoA-linked carbonyl group. This intermediate may then undergo a β-elimination to remove acetyl-CoA and restore aromaticity, yielding cinnamate as the product. Electron balance calculations (Table S15) show that the proposed activation reaction is redox-neutral: any electrons removed from benzene during activation are internally shuttled within the complex – likely via its flavin and Fe-S cofactors – and returned to the activation product.

**Figure 7.**
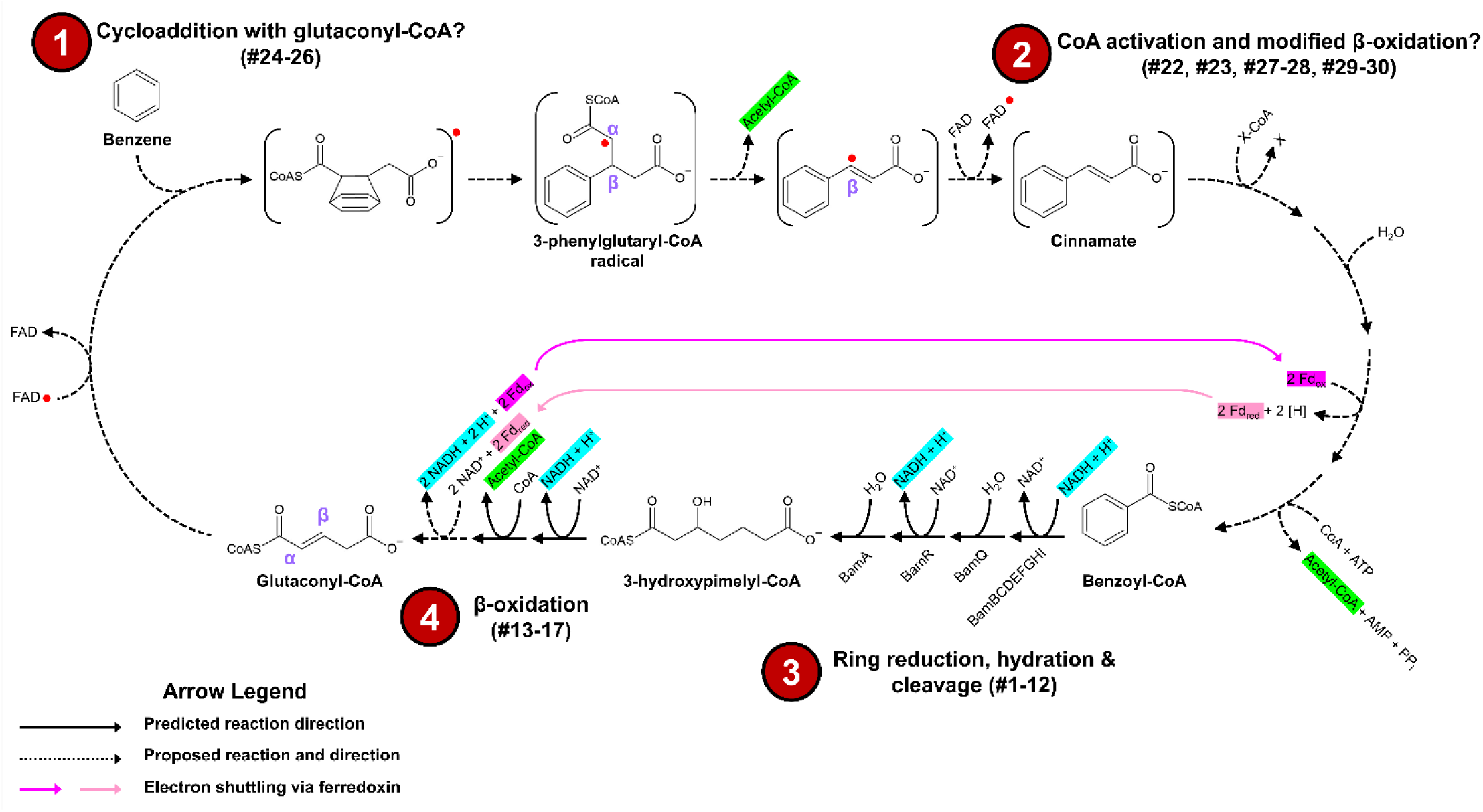
Conceptual model of anaerobic benzene degradation by ORM2a. We propose that a radical flavin-mediated cycloaddition reaction with glutaconyl-CoA is used to activate benzene in ORM2a. Radicals are denoted by red circles. This closed-loop model suggests that glutaconyl-CoA does not undergo further β-oxidation steps yielding acetyl-CoA. Rather, stoichiometric amounts of acetyl-CoA is produced from three separate lyase reactions. Ferredoxin may serve as an electron shuttle between the proposed EtfAb-glutaryl-CoA dehydrogenase complex (Proteins #15-17) and an unknown dehydrogenase during the transformation of cinnamate to benzoyl-CoA. Electron balances for each reaction step are shown in Table S15.

### Transformation of the proposed benzene activation intermediate to benzoyl-CoA

If cinnamate is the product of the initial benzene activation reaction, the most plausible next steps include coenzyme A activation followed by modified β-oxidation (85), yielding benzoyl-CoA and releasing an additional mole of acetyl-CoA (Figure 7). Stoichiometry indicates that three moles of acetyl-CoA and three moles of H_2_ or equivalent are produced per mole of benzene oxidized (86). To balance this stoichiometry, we propose that Proteins #22 and #27-28 function in appending CoA moieties to cinnamate and a downstream intermediate, enabling β-oxidation to proceed. The third mole of acetyl-CoA is released by a downstream thiolase (Protein #14) during the conversion of 3-hydroxypimelyl-CoA to glutaryl-CoA (Figures 2 and 7). The proposed metabolic pathway is also electron balanced (Table S15).

β-oxidation consists of four consecutive steps: dehydrogenation, hydration, oxidation, and thiolysis. Multiple protein prediction tools implicate Proteins #23 and #29-30 in hydration reactions (Figure 4b and Tables S13-S14). However, the “Magic” gene cluster lacks clear candidates for dehydrogenation (other than Proteins #24-26), oxidation, and thiolysis steps. It remains unclear whether these functions are carried out by enzymes encoded elsewhere in the genome or by uncharacterized proteins within the “Magic” cluster that catalyze analogous reactions.

### Closing remarks

There is little doubt that the “Magic” gene cluster in ORM2, *Pelotomaculum* candidate BPL, and specific hydrothermal vent organisms participate in anaerobic benzene degradation, and likely in its elusive, energetically difficult activation step. By contrast, the “Nanopod” gene cluster – present in ORM2a, ORM2b_MAG4, and *Deltaproteobacteria* AUK045 – likely facilitates benzene transport across the cell membrane. Given that the MAGs of most anaerobic benzene degraders surveyed were incomplete, it is possible that homologous “Nanopod” genes were missed. It is telling that the only genomes with both the “Magic” and “Nanopod” clusters belonged to strict anaerobes with apparent specialization in benzene metabolism. The location of both clusters on genomic islands (Figure 1) suggests acquisition via horizontal gene transfer. Now that these unique gene clusters have been identified, we hope the broader research community will help confirm their functions and continue the search for homologous systems in other datasets and enrichment cultures.

## MATERIALS AND METHODS

Detailed descriptions of all materials, experimental procedures, and computational analyses are provided as Supplementary Materials.

## FUNDING AND COMPETITNG INTERESTS STATEMENT

This study was funded by Genomic Application Partnership Program (GAPP) grants awarded to E.A.E. (Project IDs OGI-102 and OGI-173), which were supported by Genome Canada, Ontario Genomics, the Government of Ontario, Mitacs Canada, SiREM, Alberta Innovates, Federated Co-operatives Limited, and Imperial Oil. The authors declare no competing financial interest.

## ACKNOWLEDGEMENTS

This study was made possible by the contributions of many dedicated students and research staff over the past 15 years. Special recognition is given to Dr. Harry Beller (Lawrence Berkeley National Laboratory), Dr. Johann Heider (Philipps University of Marburg), Dr. Christine Orengo (University College London), and Dr. Joel Roca (University College London) for their invaluable guidance on characterizing proteins of unknown function.

