## Supplemental Information for "Identification of a Highly Expressed Gene Cluster Likely Coding for Benzene Activation Enzymes in a Methanogenic Enrichment Culture"

[‡]Current Address: Liven Proteins Corporation, Mississauga, Ontario, L5L 1C6, Canada

Number of Pages: 25

Materials and Methods

Number of Supplementary Texts: 1

Number of SI Figures: 11

Number of SI Tables: 13

Number of Supporting Files: 3

### TABLE OF CONTENTS

**Materials and Methods……………………………………………………………………………..4**

**Supplementary Text….…..……………………………………………………………………..10**

**Text S1**. Supplementary features of the ORM2a genome………………………………………10

**Supplementary Figures………………………………………………………………………….11**

**Figure S1.** Output results from Ori-Finder 2022……………………………………………...11

**Figure S2**. The average nucleotide identity between the closed genome of ORM2a and a draft MAG putatively belonging to ORM2b……………………………………………………...12

**Figure S6**. Maximum likelihood consensus trees showing the affiliation of Proteins #22, #43 and homologs to known and predicted AMP-binding acyl-CoA synthetase enzymes……...………..16

**Figure S7**. AlphaFold 3 predicted models of Protein #22, Protein #23, the Protein #27-28 complex, and the Protein #29-30 complex……………………………………………………….……17

**Figure S8**. Supplementary AlphaFold 3 models of the Protein #24-26 enzyme complex………...18

**Figure S9**. Maximum likelihood consensus trees showing the affiliation of Protein #25 and homologs to known and putative heterodisulfide reductase enzymes…………………………....19

**Figure S10**. Maximum likelihood consensus trees of known and putative heterodisulfide reductase enzymes……………………………………………………………………………………..20

**Figure S11**. Multiple sequence alignment of cysteine-rich motifs in Protein #25 and selected iron-sulfur homologs from methanogens and *Escherichia coli*………………………………………21

**References………………………………………………………………………………………23**

**Supplementary Tables (all tables can be found in accompanying Excel file)**

**Table S1**. Relative abundance of archaea, bacteria, and ORM2 in OR maintenance cultures sampled between 2019-2024

**Table S2**. Summary of OR metagenomes and metagenome-assembled genomes (MAGs)

**Table S3**. General features of the ORM2a MAG

**Table S4**. Complete and incomplete KEGG metabolic pathways identified in the ORM2a genome. The table is separated into a summary (**Table S4a**) and more detailed information (**Table S4b**) into the KEGG pathways identified

**Table S5**. Putative transporter genes identified in the ORM2a MAG

**Table S6**. Operons in the ORM2a MAG putatively associated with syntrophic metabolic processes and hydrogen evolution

**Table S7.** Putative genomic islands (GI) and prophage regions identified in the ORM2a MAG

**Table S8.** Putative transposase genes and pseudogenes identified in the ORM2a MAG

**Table S9.** Putative insertion sequences (IS) identified in the ORM2a MAG

**Table S10**. Proteins identified using LC-MS/MS sequencing of OR crude lysates and SDS-PAGE gel slices

**Table S11**. Computational and statistical analysis of liquid chromatography tandem mass spectrometry (LC-MS/MS) output. The table is separated by peptide (**Table S11a**) and protein (**Table S11b**) information

**Table S12**. BLASTP search results for ORM2a proteins of interest against MAG4_ORM2b and publicly available genomes in GenBank and JGI

**Table S13**. Best homology search results for ORM2a proteins coded by "Magic" and "Nanopod" gene clusters

**Table S14**. Best Gene Orthology (GO) term search results for ORM2a proteins coded by "Magic" and "Nanopod" gene clusters. The table is separated by sequence-based (**Table S14a**) and structure-based (**Table S14b**) search results

**Table S15**. Electron balances for anaerobic benzene degradation pathways catalyzed by ORM2a

**Supporting Files (available in figshare: https://doi.org/10.6084/m9.figshare.27312285.v2)**

**File S1.** FASTA file of OR proteomics database

**File S2.** AlphaFold 2 structural models of ORM2a “Magic” and “Nanopod” proteins

**File S3.** AlphaFold 3 structural models of ORM2a “Magic” enzyme complexes

### MATERIALS AND METHODS

#### Culture maintenance

The OR consortium was enriched from samples from an oil refinery in Oklahoma in 1995 (1) and has been maintained in a near-identical manner ever since. Briefly, cultures are grown in a defined pre-reduced anaerobic mineral medium (2, 3) and are fed benzene (~5-40 mg/L) once every 4-6 weeks. The OR consortium has been subcultured many times, resulting in the emergence of five distinct culture lineages (4, 5). The "OR-b" lineage, illustrated in Figure S3, is the most well characterized with a doubling time about 30 days (3-10) and was surveyed in this study. A large scale (>100 liters) bioaugmentation culture known as DGG-B (3, 9) was derived from the OR-b lineage. Though the OR consortium has been amended with other substrates including benzoate and toluene, only benzene has been shown to support the growth of ORM2a and ORM2b (6, 11).

#### Sequencing, assembly, and annotation of OR metagenomes and MAGs

Between 2010-2017, the metagenomes of three OR-b subcultures designated OR-b1A, DGG1A, and DGG-0 were sequenced. The OR-b1A (2010) metagenome was sequenced using Illumina paired-end technology (2×100 bp) and was partially assembled in ABySS v.1.3.2 (12) with contigs binned using varying kmer lengths. Details are available in Chapter 5 of Devine's PhD thesis (6). Metagenomic sequencing and assembly of DGG1A (2016) and DGG-0 (2017) was described as part of a Microbiology Resource Announcement by Toth et al. (8). Briefly, hybrid assembly of short-read (Illumina paired-end) and long-read (PacBio) shotgun sequences was utilized to produce long, high quality contigs. Following binning, three draft MAGs including ORM2a were refined into closed (completed) genomes (8). The correct assembly of each closed genome was verified by read mapping, and the average sequence depth for the ORM2a genome was 1,384× (8, 13).

Taxonomy was assigned to all draft and complete MAGs using release 214 of the Genome Taxonomy Database (GTDB) and GTDB tool kit (v.2.3.2) using the classify_wf workflow (8, 14, 15). In this study, the ORM2a genome (CP113000.1) and a medium-quality MAG putatively belonging to ORM2b, (ORM2b_MAG4, 3,214,903 bp across 378 contigs) were reclassified using release 220 of GTDB and GTDB-Tk v.2.4.0 (discussed later). Seventy-one medium-to-high quality MAGs including ORM2b_MAG4 were retained. All metagenomic assemblies and complete MAGs were submitted to NCBI and/or IMG for automated gene calling and functional annotation. Incomplete (noncircular) MAGs were deposited to figshare (13) and in this study were annotated using RASTtk (16) in March, 2022. Data availability including accession numbers and all functional annotation pipelines used is provided in Table S1.

**Analysis of the ORM2a genome**

To assess the metabolic potential of the ORM2a genome, coding sequences predicted by the NCBI Prokaryotic Genome Annotation Pipeline were uploaded to BlastKOALA for automatic KEGG Orthology (KO) assignment (17), then organized into metabolic pathways using the Reconstruct tool within KEGG Mapper (18, 19). Next, the mummer2circos package in GitHub (https://github.com/metagenlab/mummer2circos) (20) was used to generate a circular plot of the ORM2a genome and to compare its homology with ORM2b_MAG4 (Figure 1). Two-way average nucleotide identity (ANI) analysis of both genomes was performed using an online calculator (http://enve-omics.ce.gatech.edu/ani/). Ori-Finder 2022 (21) was used to identify the genomes' origin of replication (*oriC*). IslandViewer 4 (22) and PHASTER (23) were used to identify genomic islands and prophage sequences, respectively. Functional annotations and hidden Markov model assignments (from NCBI and IMG) were used to identify transposase genes. Putative insertion sequences were identified using ISfinder (24). Operon-mapper (25) was used to identify operons in the ORM2a genome.

**Protein extraction and LC-MS/MS analysis**

Between 2010-2018, three proteomics experiments were performed on three OR-b lineage subcultures (OR-b1C, OR-b, and OR-b1A; see Figure S3). Culture information, including culture volumes extracted and benzene degradation rates at the time of sampling, are provided in Table 1. To avoid oxygen contamination, all containers used were incubated in an anaerobic chamber (supplied with a gas mix of 10% $H_2$, 10% $CO_2$ and 80% $N_2$) for >24 hours prior to use, and all steps were performed at 4 °C unless otherwise specified.

Protein extractions for Experiment #1 (OR-b1C, 2010) and Experiment #2 (OR-b, 2011) were conducted by Devine (6). Cell pellets from 150-200 mL of culture were harvested by centrifugation (8,000 x g, 20 min), flash-frozen using liquid nitrogen, and stored at -80°C until further processing. On the day of protein extraction, thawed cells were resuspended in a lysis buffer solution (100 mM Tris-HCl [pH 8.0], 5% w/v glycerol, 10 mM EDTA, 1 mM PMSF [phenylmethylsulfonyl fluoride], and 5 mM dithiothreitol [DTT]) and sonicated at 23-30W for 10 min in pulse mode (1 s ON/1 s OFF for 1 min, 1 min OFF, repeat). Proteins in the crude lysate were recovered using phenol [pH 8.0] and precipitated with 100 mM ammonium acetate in methanol. After centrifuging (10,000 x g, 20 min) and discarding the supernatant, the protein pellets were washed with acetone, resuspended in a 6 M urea solution (pH 8.0), then subjected to an 18-24 hr in-solution digestion with porcine trypsin (19). The digestion was stopped using a solution of 2% trifluoroacetic acid (TFA) and 20% acetonitrile. The resulting peptides were purified using PepClean™ Spin Columns (Pierce Biotechnology) according to the manufacturer's procedure.

In Experiment #3 (OR-b1A, 2018), cell pellets from two 50 mL volumes of culture (10,000 x g, 15 min) were suspended in an anoxic sodium dodecyl sulfate (SDS)-containing lysis buffer (50 mM Tris-HCl [pH 7.6], 2% w/v SDS, and 50 mM DDT) and sonicated at 23-30W for 10 min in pulse mode (1 s ON/1 s OFF for 1 min, 1 min OFF, repeat). Proteins from one crude lysate (Experiment #3a) were purified using a Microcon-30kDa centrifugal filter unit (Millipore) as outlined by Wiśniewski et al. (26), then subjected to an 18-24 hr in-solution digestion with porcine trypsin at 37 °C (6). The second crude lysate (Experiment #3b) was mixed with 2× SDS gel-loading buffer (50 mM Tris-HCl [pH 6.8], 10% SDS [sodium dodecyl sulfate], 40% glycerol, 3 mM bromophenol blue, and 500 mM DDT]) and separated on a 15% SDS-PAGE gel run for 45 min at 200 V. Visualization of the resulting gel revealed a long protein smear with no distinct bands (Figure S2), which was sliced into five equal sections then destained and digested with trypsin as described in Shevchenko et al. (27).

LC-MS/MS analysis of Experiments #1 and #2 peptides (6) was performed at the SickKids SPARC BioCentre using an LTQ Orbitrap mass spectrometer (Thermo Fisher Scientific). Run parameters are summarized in Tang et al. (28). LC-MS/MS analysis of Experiment #3 peptides was conducted at the University of Toronto BioZone Mass Spectrometry Facility. Briefly, 5 µL liquid samples of peptides were separated on a 15cm PicoTip Emitter (New Objective) packed with ReproSil-Pur C18-AQ 3 µm resin (Dr. Maisch GmbH). The flow rate was set to 250 nL/min and the eluents used were (A) water containing 0.1% formic acid, and (B) acetonitrile containing 0.1% formic acid. The gradient started at 0% B, followed by a linear gradient to 10% B over 5 min, a linear gradient to 40% B over 88 min, a linear gradient to 95% B over 2 min, a hold at 95% B for 10 min, a linear gradient to 0% B over 1 min, and a final hold of 0% B for 14 min (total runtime of 120 min). MS detection was conducted using a Q-Exactive Orbitrap mass spectrometer

(Thermo Fisher Scientific) equipped with a nanoelectrospray ionization probe operating in positive ionization mode, with a spray voltage of 2.8 kV, capillary temperature of 275°C, and S-lens radio frequency level of 55. Full MS data was acquired with an m/z range of 400-2,000, mass resolution of 70,000, automatic gain control (AGC) target of 1.0E+06, and a maximum injection time of 30 milliseconds. MS2 was gathered using a data dependant TOP10 approach with a mass resolution of 17,500, AGC target of 5.0E+04, maximum injection time of 50 ms.

**Protein identification**

Raw mass spectrometry data from all three experiments were converted into mzXML using MSconvert (29) then uploaded into Version 2020.11.12.1 of X! Tandem (The GPM, thegpm.org) for peptide and protein discovery. Spectra were screened against a large library of protein sequences (1,595,050 entries) with a mass tolerance of <0.4 Da for the fragment ions and <20 ppm for the precursor ion. The library, provided in File S1 and overviewed in Table S1, contained 1,106,523 translated gene sequences called by IMG (three metagenomic assemblies and 3 closed MAGs), NCBI (3 closed MAGs), and RAST-tk (67 draft MAGs). The database also included 488,411 reverse-translated (decoy) protein sequences from the DGG0 metagenome to calculate a false discovery rate, and 116 entries from common human/laboratory contaminants. Peptide and protein identifications were accepted if they could be established at greater than 95% and 99% probability, respectively (30, 31). Proteins sharing peptide evidence were grouped into clusters.

**Functional and structural prediction of ORM2a proteins detected in high abundances**

Protein identity was verified using protein BLAST searches against all other OR metagenomic assemblies and MAGs, as well as reference (meta)genomic sequences in NCBI (nr), JGI (IMG/M), and UniProt (UniProt KB reference proteomes + Swiss-Prot). Next, the sequence-based tool MotifFinder (32) was used to identify conserved domains in each protein. Tertiary protein structures were predicted using the AlphaFold2.ipynb tool within Colab (33, 34), then compared to homologous structures and protein domains within the Dali server's Protein Data Bank (35) and DeepFRI (36). All AlphaFold 2 protein structural models generated are provided in File S2. Additionally, AlphaFold 3 (37) was used to predict enzyme complex structures for select proteins of interest. Maximum likelihood trees were constructed in RAxML v.8.2.1.1 within Geneious v.8.1.9 (38, 39) to corroborate predicted protein functions where possible. Protein sequences were also sent to the laboratory of Dr. Christine Orengo (University College London) for independent functional analysis using CATH v.4.4 (40) and PROST (41).

**Phylogenomic analyses**

Phylogenomic analysis of the closed ORM2a genome and ORM2b_MAG4 was performed in v.1.8.8 of GToTree (42). Briefly, we retrieved all representative genomes from c__UBA8473 and c__WTBG01 from GTDB v.220, as well as closely related genomes from selected *Desulfobacterota*, then extracted target genes using the Proteobacteria.hmm single copy gene set (119 genes) within GToTree. *In silico* translated protein sequences were aligned using MUSCLE (43) and concatenated. A maximum likelihood tree was constructed from the concatenated alignment using IQ-TREE v.2.3.6 (44) with the LG+F+I+R5 substitution model, inferred as best model by IQ-TREE, and 100 bootstraps. For comparative purposes, a second maximum likelihood tree was constructed using 16S rRNA gene sequences extracted from each representative GTDB genome (if available) and from clone sequences of known/predicted anaerobic benzene degraders. *Desulfocapsa sulflexigens* DSM 10523 (GCA_000341395.1) was included as an outgroup.

**Text S1**. Supplementary features of the ORM2a genome

Twelve possible genomic islands were identified in the ORM2a genome, of which two are likely prophage sequences and 9 contain putative transposase genes (Figure 1 and Table S7). An additional 47 transposase genes and 63 insertion sequences-like structures were scattered across the genome (Figure 1 and Tables S8-S9), hinting that the ORM2a genome has been shaped by numerous rearrangement events and may explain the absence of a definitive terminus region. A similarly high transposase content was found in the closed genome of "*Ca*. Nealsonbacteria" DGGOD1a, a predicted necromass recycler (5). No CRISPR elements, virulence factors, or antimicrobial resistance genes were identified. Genes coding for type IV pili and a Che-type chemotaxis system may be involved in bacterial motility.

Only 29 complete anabolic pathways were identified in the ORM2a genome, including biosynthesis pathways for 7 amino acids and 4 vitamins (Table S4a). In contrast, 197 genes with putative transport functions were identified (Table S5) – including transporters for vitamin $B_{12}$ and undefined amino acids – hinting that ORM2a imports most of its nutrients from extracellular sources, perhaps from polymeric substances or from direct exchanges with other microorganisms as theorized in recent microscopy and genomic investigations of the OR culture (5, 10). Similar transporter gene counts were recovered from the genome of *Syntrophus acidotrophicus* SB, a fermentative benzoate-degrading anaerobe whose central and peripheral metabolic pathways are also largely incomplete (45, 46).

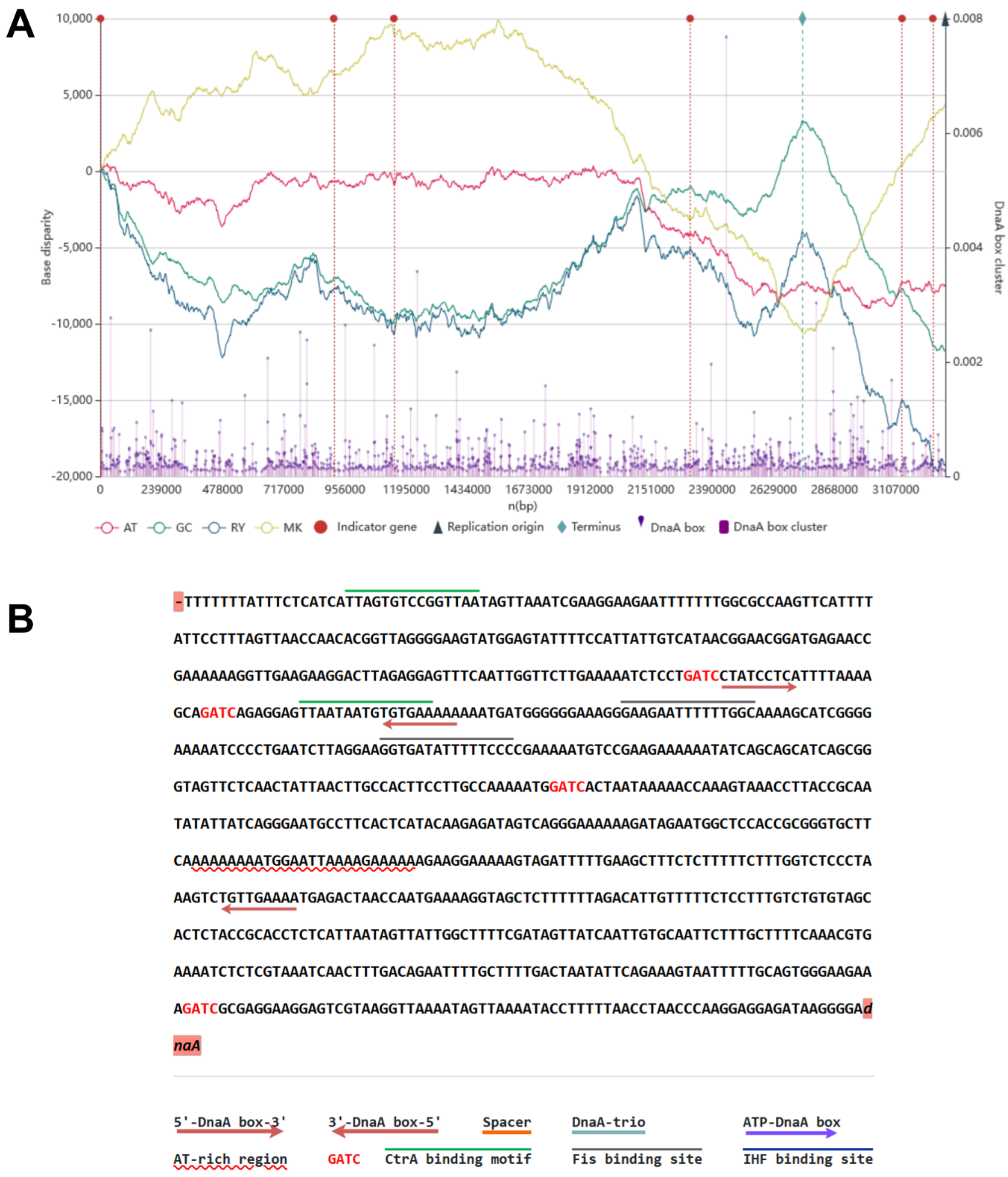

**Figure S1**. Output results from Ori-Finder 2022 (21). The top panel (a) shows RY, MK, AT and GC disparity curves of the ORM2a genome, the location of 5 indicator genes (*dnaA*, 790…2,136 nt; *hemE*, 911,117…912,208 nt; *dnaN*, 1,146,408…2,304,626 nt; *mnmG*, 3,131,767…3,133,644 nt, *hemB*, 3,251,271…3,252,257 nt), the predicted replication origin (*oriC*), the maximum of GC disparity (2,742,827 nt), DnaA boxes (i.e., DnaA binding sites), and DnaA box clusters. The bottom panel (b) shows the *oriC* sequence including specific binding motifs. The *dnaA* gene is located immediately downstream of *oriC*.

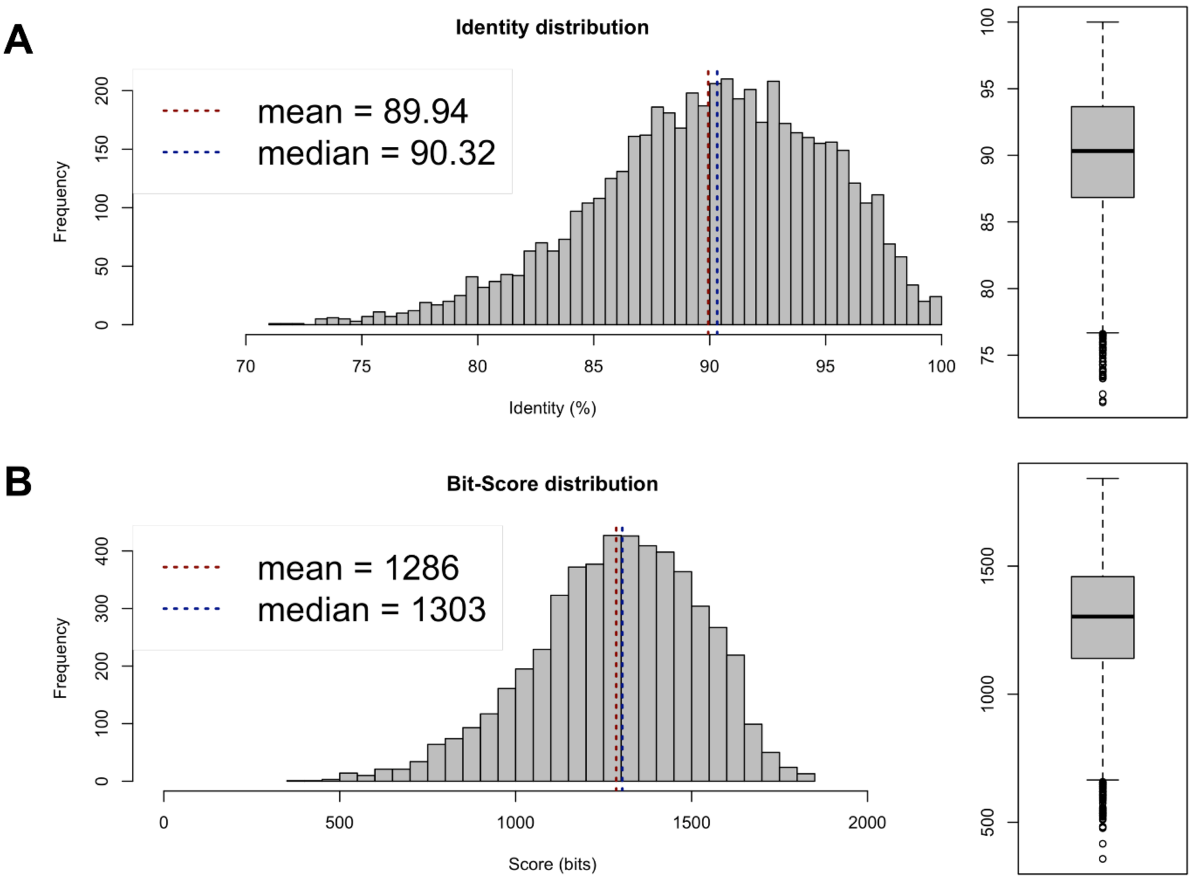

**Figure S2**. The average nucleotide identity (ANI) between the closed genome of ORM2a (CP113000.1) and a draft MAG putatively belonging to ORM2b (ORM2b_MAG4). Panel a) shows the best reciprocal BLAST hits (two-way ANI) between each genome pair where aligned coding regions share ≥70% identity. Panel b) shows the statistical significance (bit-score) of each alignment. Figures and ANI values were generated using the ANI calculator developed by the Kostas lab (http://enve-omics.ce.gatech.edu/ani/).

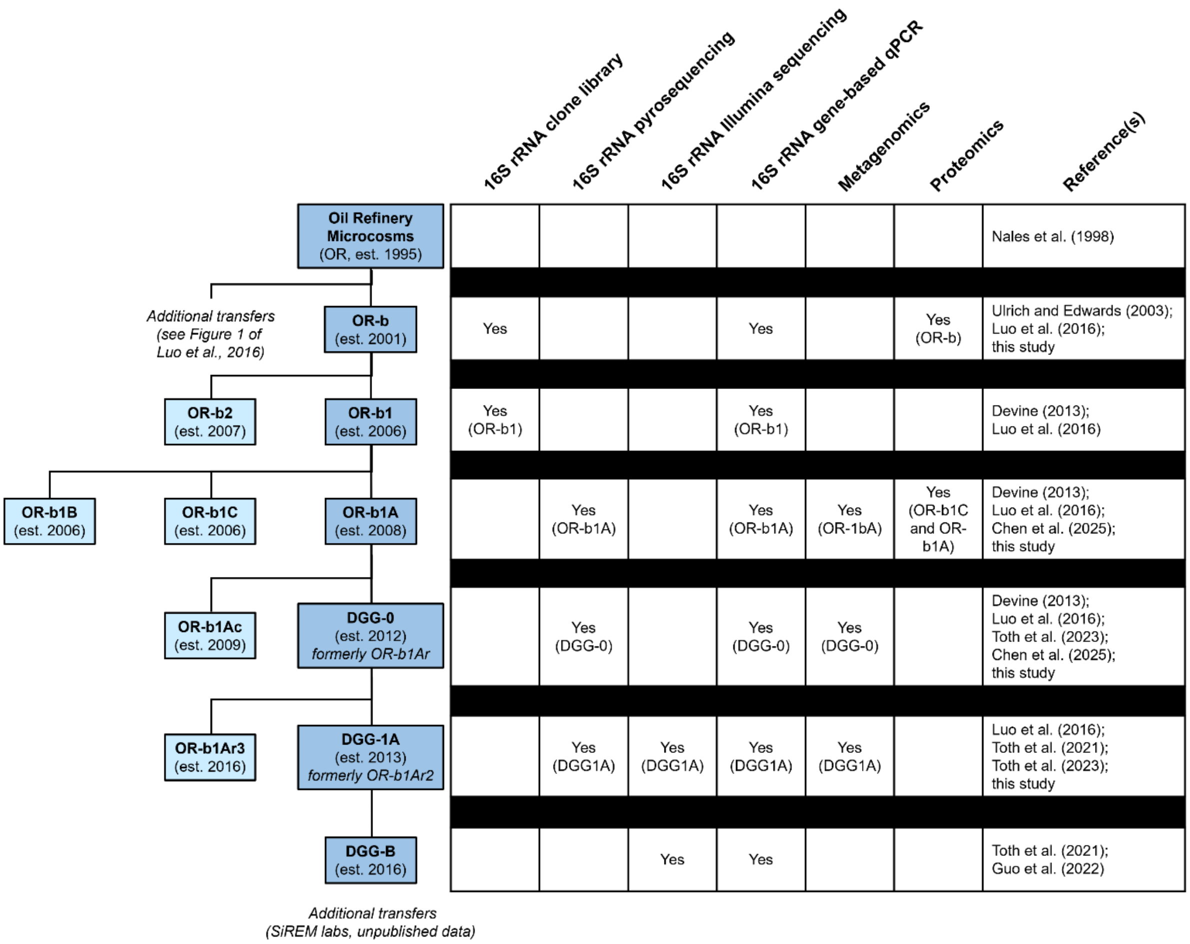

**Figure S3**. Subculturing history of the methanogenic OR consortium's "OR-b" lineage, including molecular analyses performed to date.

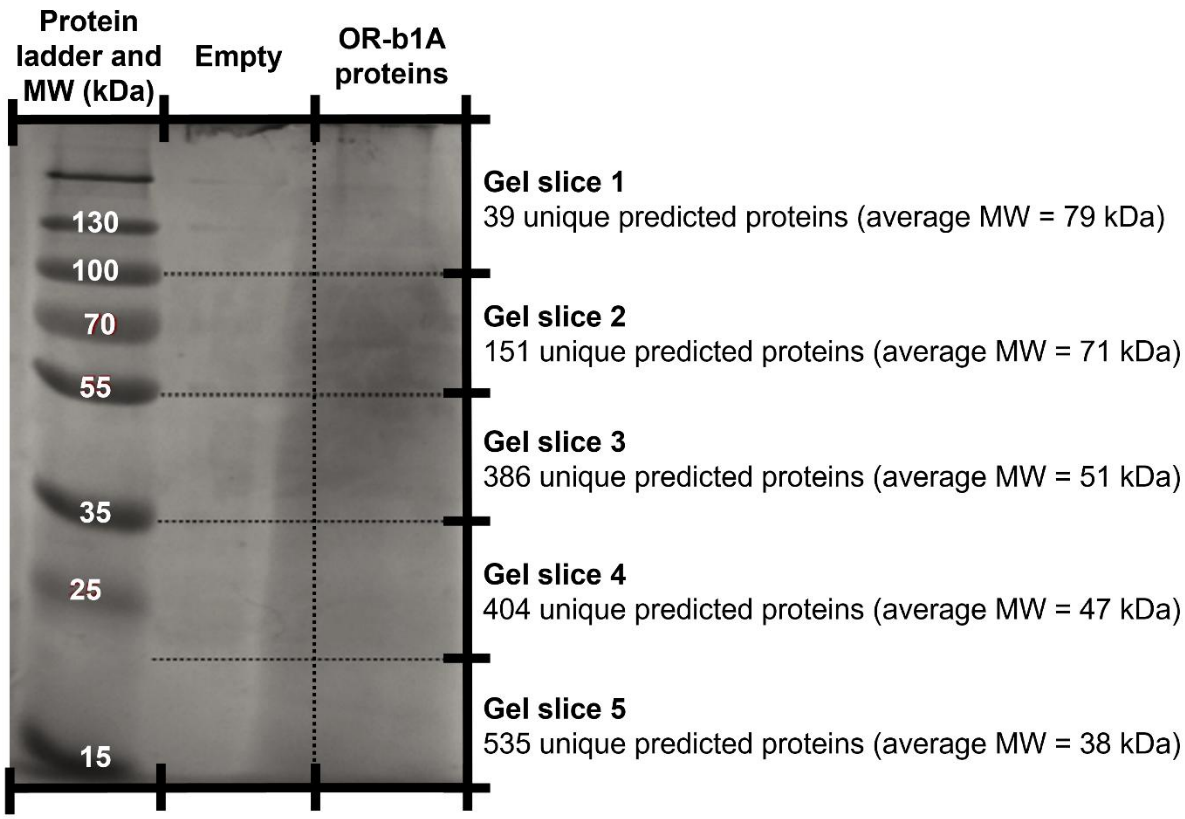

**Figure S3**. Polyacrylamide gel electrophoresis (SDS-PAGE) of proteins extracted from Experiment #3b. Proteins from a commercial ladder (left-most lane) and the crude extract from OR-b1A (right-most lane) are shown. Dotted lines show the approximate locations where the OR-b1A protein band was sliced in preparation for in-gel trypsin digestion and LC-MS/MS sequencing (Experiment #3b). MW = molecular weight.

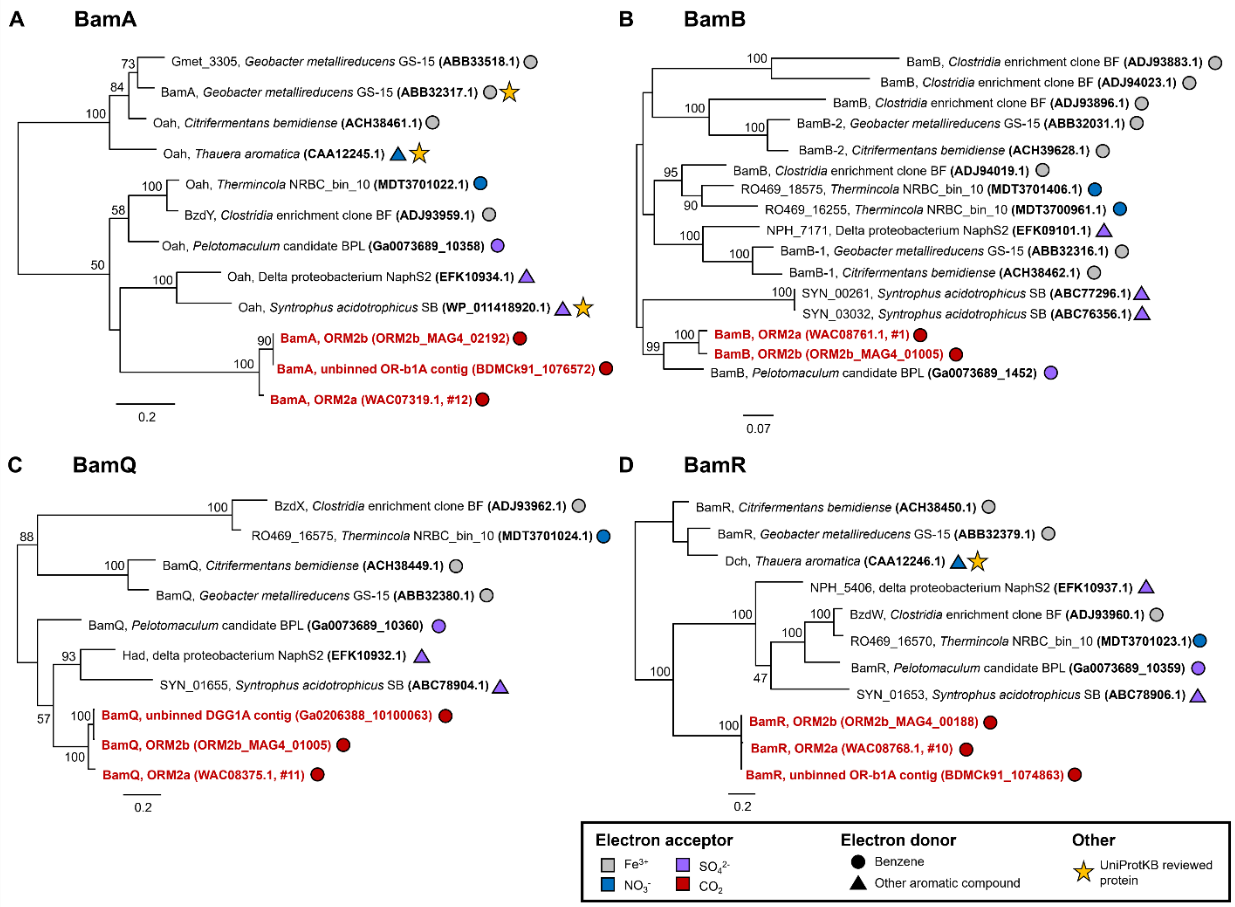

**Figure S5**. Maximum likelihood consensus trees showing the affiliation of predicted OR consortium *bam* gene products – 6-oxo-cyclohex-1-ene-carbonyl-CoA hydrolase (panel a), benzoyl-CoA reductase subunit b (panel b), 6-hydroxycyclohex-1-ene-1-carbonyl-CoA dehydrogenase (panel c), and cyclohexa-1,5-diene-1-carbonyl-CoA hydratase (panel d) – to reference protein sequences from select anaerobic aromatic degraders. Sequences highlighted in red are from this study. Proteins marked with stars have some experimental evidence of protein function and have been reviewed by UniProtKB. Bootstrap values < 50% are not shown. GenBank accession numbers are provided for most proteins; IMG accession numbers are provided for *Pelotomaculum* candidate BPL and the OR-b1A metagenome. Amino acid sequences for ORM2b_MAG4 are available in Table S12. Multiple sequence alignment statistics for each protein are also available in Table S12.

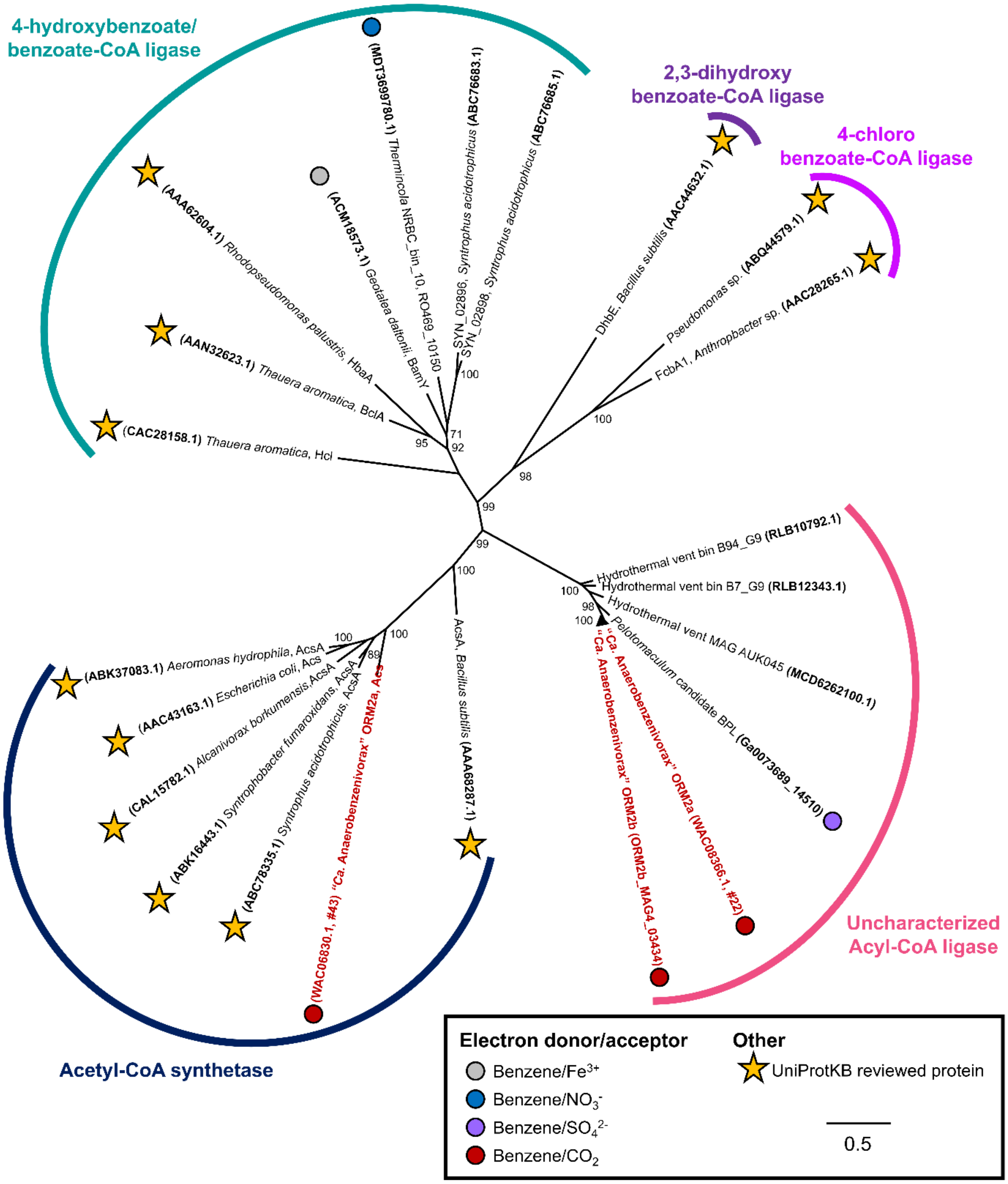

**Figure S6**. Maximum likelihood consensus trees showing the affiliation of Proteins #22, #43 and homologs to known and predicted AMP-binding acyl-CoA synthetase enzymes. Proteins marked with stars have some experimental evidence of protein function and have been reviewed by UniProtKB. Bootstrap values < 70% are not shown. Multiple sequence alignment statistics for Proteins #22 and #43 are also available in Table S12.

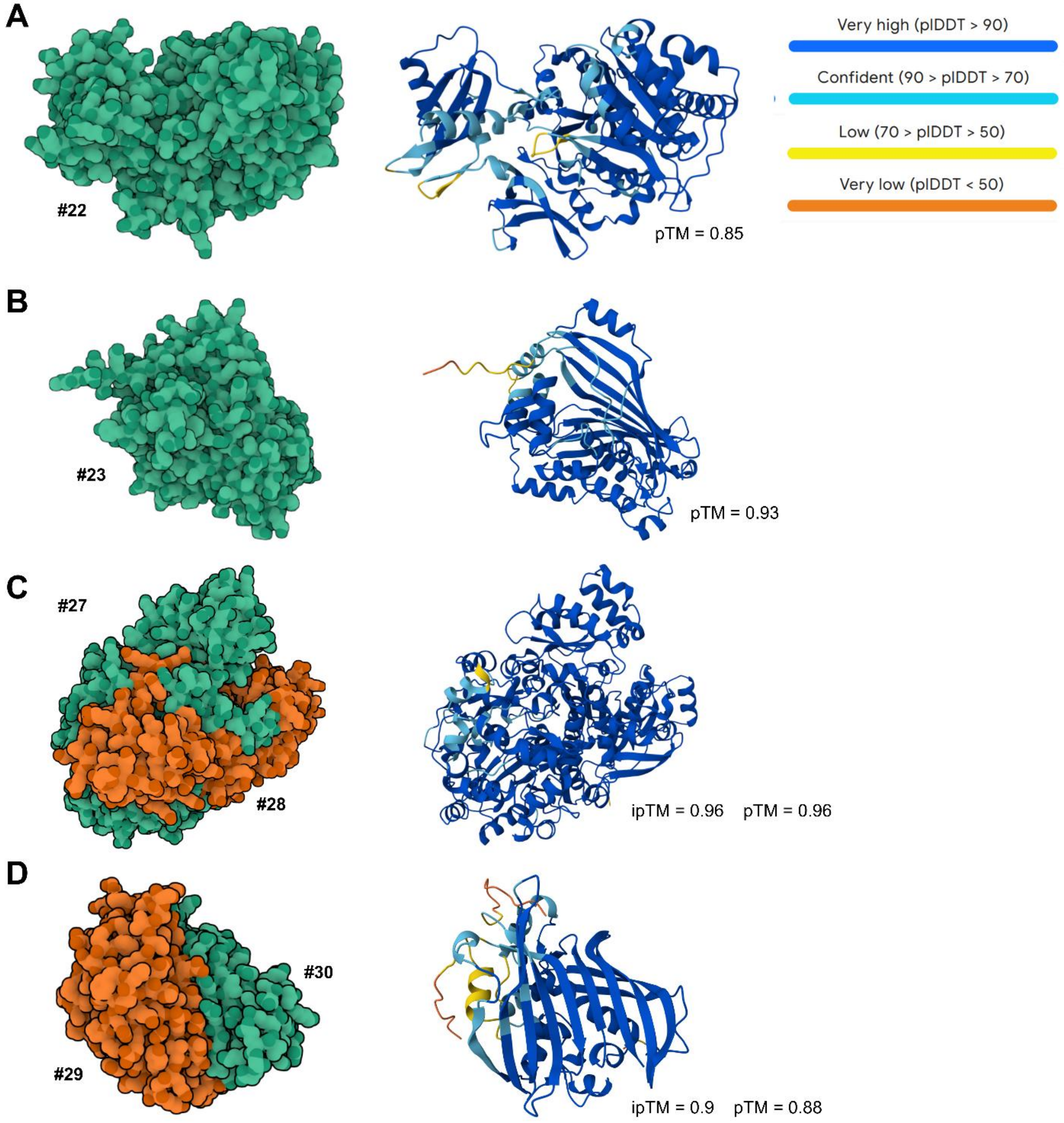

**Figure S7**. AlphaFold 3 predicted models of a) Protein #22, b) Protein #23, c) the Protein #27-28 complex, and d) the Protein #29-30 complex. Surface and ribbon models of each structure are shown. The colour outputs of each ribbon model depict predicted Local Distance Difference Test (pIDDT) confidence estimate of each atom on a 0-100 scale, where a higher value indicates higher confidence. The predicted template modeling (pTM) score and the interface predicted template modeling (ipTM), which estimates the folding accuracy of predicted structures and protein complexes, respectively, is also shown on a 0-1 scale, where a higher value indicates higher confidence.

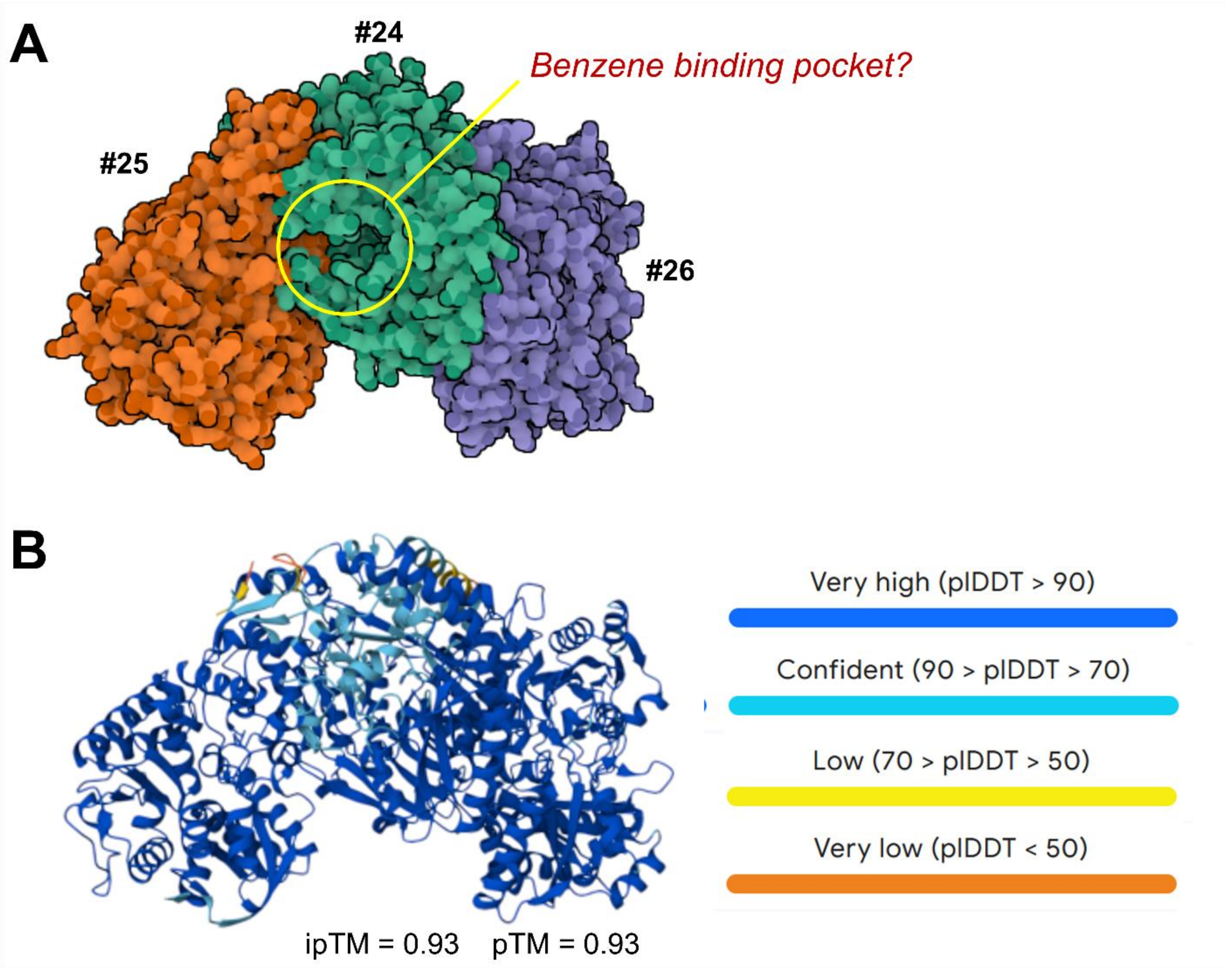

**Figure S8**. Supplementary AlphaFold 3 models of the Protein #24-26 enzyme complex. Surface (panel a) and ribbon (panel b) models of each structure are shown. A putative binding pocket is shown in the surface model of Protein #24. The colour outputs of the ribbon model depicts the predicted Local Distance Difference Test (pIDDT) confidence estimate of each atom on a 0-100 scale, where a higher value indicates higher confidence. The predicted template modeling (pTM) score and the interface predicted template modeling (ipTM), which estimates the folding accuracy of predicted structures and protein complexes, respectively, is also shown on a 0-1 scale, where a higher value indicates higher confidence.

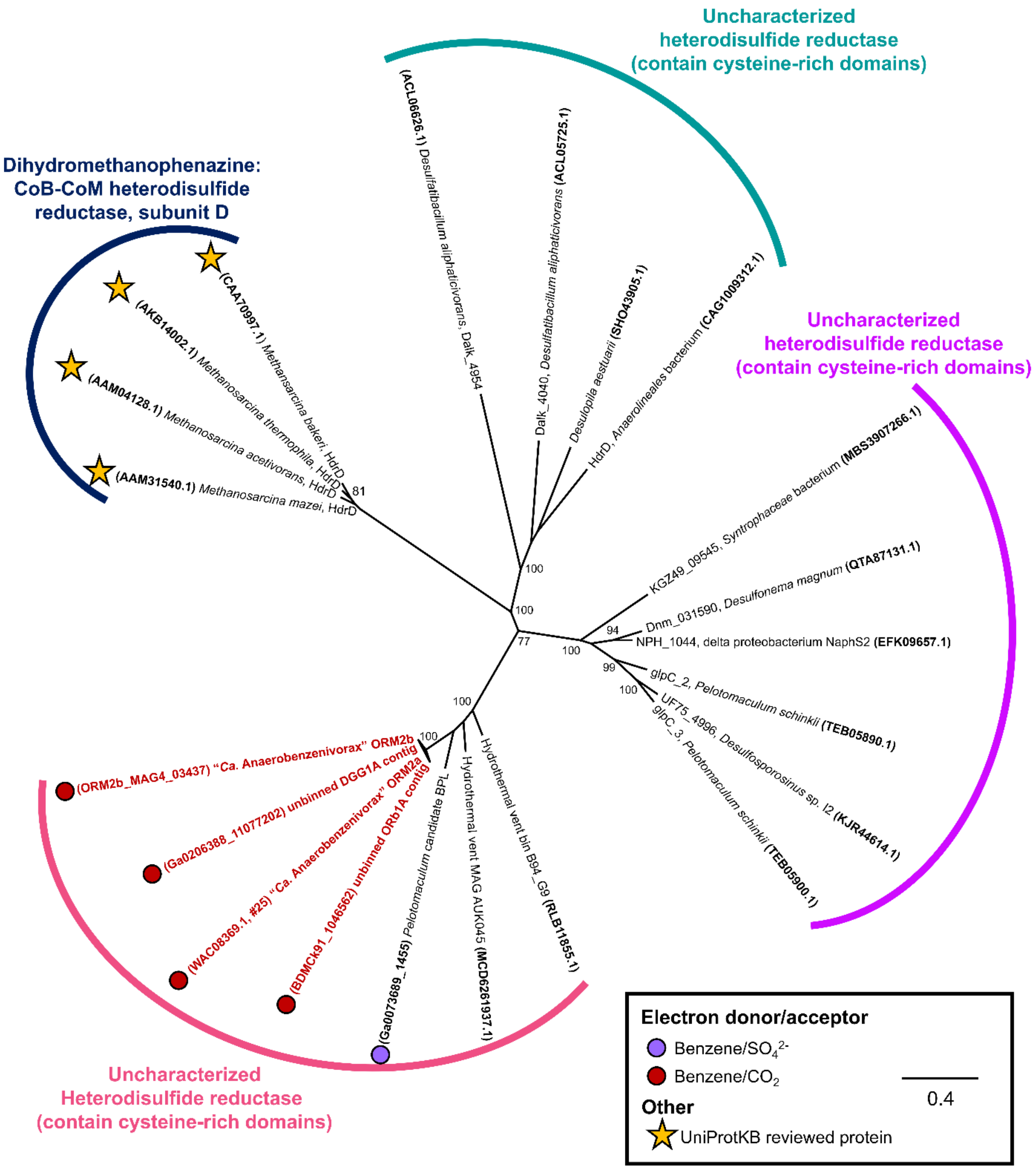

**Figure S9**. Maximum likelihood consensus trees showing the affiliation of Protein #25 and homologs to known and putative heterodisulfide reductase enzymes. Two proteins that mapped to unbinned OR contigs are also shown, as they share close homology (95-99% identity) with Protein #25. Proteins marked with stars have some experimental evidence of protein function and have been reviewed by UniProtKB. Bootstrap values < 70% are not shown. Multiple sequence alignment statistics for most proteins are available in Table S12.

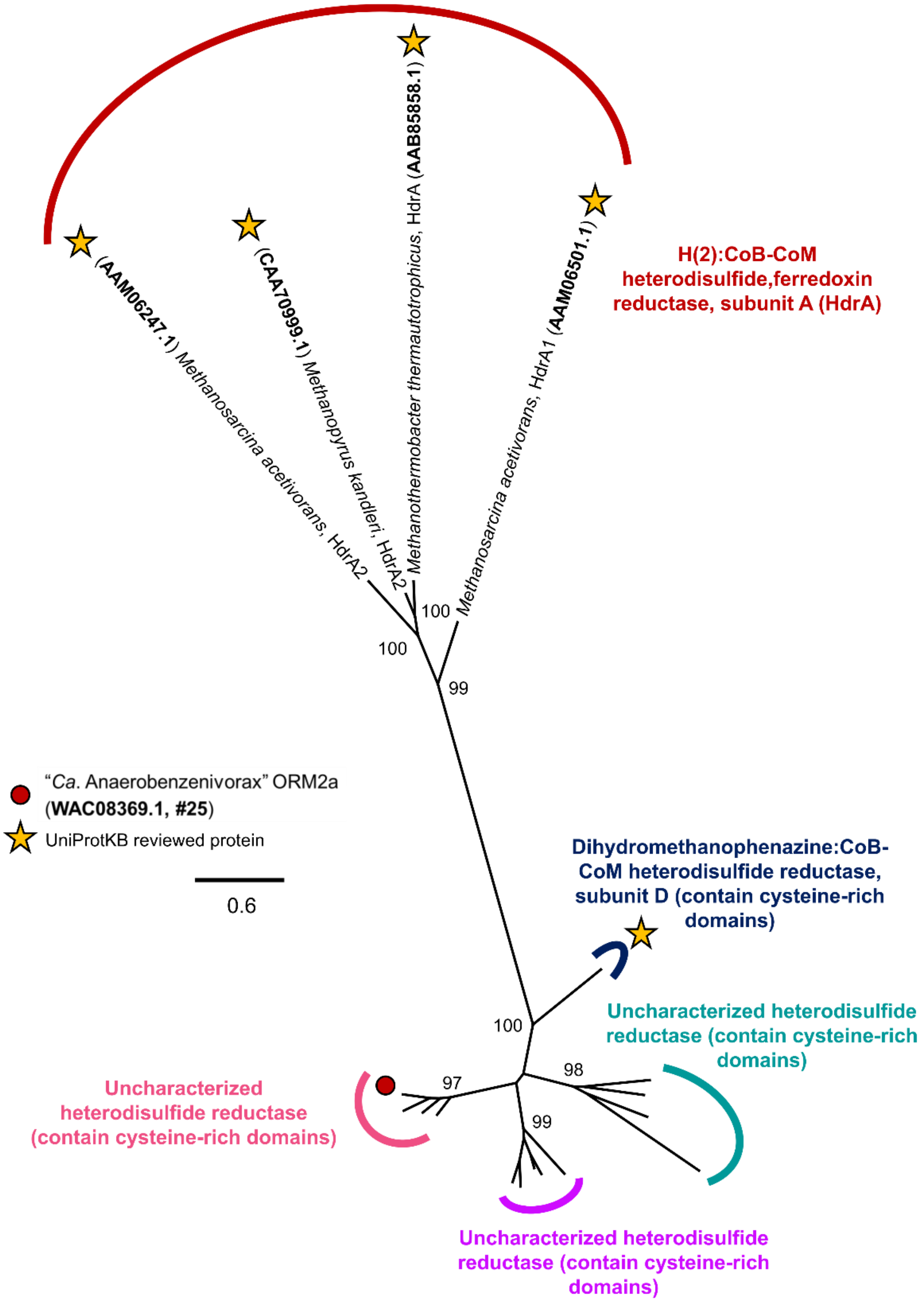

**Figure S10**. Maximum likelihood consensus trees of known and putative heterodisulfide reductase enzymes. This tree contains the same sequences and clustering pattern as Figure S9 but includes four HdrA sequences from methanogenic archaea. Proteins marked with stars have some experimental evidence of protein function and have been reviewed by UniProtKB. Bootstrap values < 70% are not shown.

**A**

```
MtHdrB    ------------------------------------------------------------    0
MhHdrB    ------------------------------------------------------------    0
#25       ----MTNTFKLDHNLDNYEWESRCCMGCAMCKHGDWIHVASDHNFSWICPEWQWG----K    52
GlpC      ----------------MNDTSFENCIKCTVCTTA--------------CPVSRVNPGYPG    30
GlcF      MQTQLTEEMRQNARALEADSILRACVHCGFCTAT--------------CPTYQLL----G    42
YkgE      ------------------------------------------------------------    0

MtHdrB    ------------------------------------------------------------    0
MhHdrB    ------------------------------------------------------------    0
#25       FDNWGACGRTRIINSLLYGDLEYTDTLAQEAAFRCFSCGGCDVSDKRNLDLEILMMNQSL    112
GlpC      PKQAGPDGERLRLKD---------GALYDEALKYCINCKRCEVACPSDVKIGDIIQRAR-    80
GlcF      DELDGPRGRIYLIKQVLEGN--EVTLKTQEHLDRCLTCRNCETTCPSGVRYHNLLDIGR-    99
YkgE      ------------------------------------------------------------    0
```

**B**

```
MtHdrB    ---------------------------------------MKYAFFLGCIMPNRYAGVEAA    21
MhHdrB    --------------------------------------MHEYAFFLGCIAPNRYPGCEAS    22
#25       ---------LERTGNYYGLNQKDRKNWVAS--DIKVEKKADTLFFVGCWASFKDNAVAQA    181
GlpC      LLDAALKIDHRRTLPKYSF--GTFRRWYRSVAAQQAQYKDQVAFFHGCFVNYNHPQLGKD    181
GlcF      --------QVRAKLPAETV---------KAKPRPPLRHKRRVLMLEGCAQPTLSPNTNAA    188
YkgE      ---------------------------------------MNVNFFVTCIGDALKSRMARD    21

MtHdrB    TRTVMEKLGVELVDMTGASCCPAPGVFGSFDQKTWLTLAARNL---C-IAEEMGVDIVTV    77
MhHdrB    AIKTSEKVGIKLLPLKGASCCPAPGAFGSIDLNVWYAMAARNL---V-LAEEMKKDIALI    78
#25       TARVLNKAQVPFMMLDNETCCGNLQYQTGHLEKF-KSAARFNLEKIRA---TGAKKVVAS    237
GlpC      LIKVLNAMGTGVQLLSKEKCCGVPLIANGFTDKA-RKQAITNVESIREAVGVKGIPVIAT    240
GlcF      TARVLDRLGISVMPANEAGCCGAVDYHLNAQEKG-LARARNNIDAWWPAIEAGAEAILQT    247
YkgE      SVLLLEKLGCRVNFPEKQGCCGQPAINSGYIKEA-IP----GMKNLIAALEDNDDPIISP    76

MtHdrB    CNGCYGSLF-EAAHLLHDNKEALNFVNEKLDKVGKEYKGNVKVRHFAELIYNDI--GVDK    134
MhHdrB    CNGCYKSIW-EVNHILKHNDELRDNVNEVLAEIDMQFKGTIDVWHLAELYYDDKVCGVQK    137
#25       CAECYRNLKVEYPKVLDFATKDLGFEVLH----------S------AELVDGLVKEGKLK    281
GlpC      SSTCTFALRDEYPEVLNVDNKGLRDHIEL----------A------TRWLWRKLDEGKTL    284
GlcF      ASGCGAFVK-EYGQMLKNDALYADKARQV----------SELAVDLVELLREE---PLEK    293
YkgE      AGSCTYAVK-SYPTYLADEPEWASRAAKV----------AARMQDLTSFIVNKL--GVVD    123
```

**C**

```
MtHdrB    IAEKVERPLN-INVGVHYGCHFLKPT-----------------------------DVKHL    164
MhHdrB    IKDSVTTPLSGAKVAAHYGCHLMKPK-----------------------------KERHF    168
#25       AKG----KMS-MKATYHDPCSLGRLSEPWIPWEGIRTGGDISNSGWGQIHPRREFRRGQF    336
GlpC      PLK----PLP-LKVVYHTPCHMEK-M----------------------------------    304
GlcF      LAI----RGD-KKLAFHCPCTLQHAQ----------------------------------    314
YkgE      VGA----SLQ-GRAVYHPSCSLARKL----------------------------------    144

MtHdrB    GSAERPVMLDEIVEATGAKSVPYADKMMCCGAGGGVRARELELSLDM---TNEKIENMIK    221
MhHdrB    GDTENPMWFEELIGALGAEPIQYRNKMQCCGAGGGVRGYDIVHALDI---TNEKLINIQE    225
#25       GCYEAPRN--LIKAIPGVELVEMLRNHSNSYHSGEGGGVYEAYPETAQFASDWRVKEAGL    394
GlpC      GWTLYTLE--LLRNIPGLELTVL--DSQCCGIAGTYGFKKENYPTSQ-AIGAPLFRQIEE    359
GlcF      KLNGEVEK--V-LLRLGFTLTDVPDSHLCCGSAGTYALTHPDLARQL---RDNKMNALES    368
YkgE      GVKDEPLT--LLKNVRGLELLTFAEQDTCCGFGGTFSVKMAEISGEM---VKEKVAHLME    199

MtHdrB    AGADCTVNVCPFCHLQFDRGQIEIKEKFGKEYNFPVLHLSQLLGLAMGMDPKDLALSVHQ    281
MhHdrB    AGADAITELCPFCQLQFDRGQIEIKEKFGDVYNIPVLHYNELLGLAQGMSPQDLALDLHA    285
#25       TGASAIITGCVHSKFMLNGALARVQNG-----INKVSHVIELVDEAYNK-----------    438
GlpC      SGADLVVTDCETCKWQIEMSTS-----------LRCEHPITLLAQALA------------    396
GlcF      GKPEMIVTANIGCQTHLASAGRT-----------SVRHWIEIVEQALEKE----------    407
YkgE      VRPEYLIGADVSCLLNISGRLQREG-Q-----KVKVMHIAEVLM---SR-----------    239
```

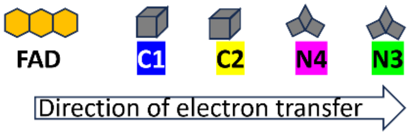

**Figure S11 (above)**. Multiple sequence alignment of cysteine-rich motifs in Protein #25 and selected iron-sulfur homologs from methanogens and *Escherichia coli*, specifically HdrB subunits from *Methanothermococcus thermolithotrophicus* (MtHdrB, PDB: 5ODQ) and *Methanospirillum hungatei* (MhHdrB, PDB: 7BKE), *E. coli* glycerol-3-phosphate dehydrogenase (GlpC, UniProt: P0A996), *E. coli* glycolate dehydrogenase (GlcF, P52074), and *E. coli* L-lactate utilization protein (YkgE, P77252). Shown in top panel (a), the N-terminal regions of GlpC and GlcF are predicted to bind two cubane [4Fe-4S] clusters via an 8-Cys motif (highlighted blue and yellow), of which 7-Cys are conserved in Protein #25 (panel a). In the center panel (b), MtHrdB and MhHrdB bind the first non-cubane [4Fe-4S] cluster via a 5-Cys motif (green), of which 4- or 5-Cys are conserved in the bacterial proteins. In the bottom panel (c), MtHrdB and MhHrdB bind the second non-cubane [4Fe-4S] cluster via a 5-Cys motif (pink), of which 4- or 5-Cys are conserved in the E. coli proteins, and only 2-Cys are conserved in Protein #25. The schematic shows the predicted electron flow from FAD to the cubane and non-cubane [4Fe-4S] clusters.
